# Short-term oxaliplatin exposure according to established hyperthermic intraperitoneal chemotherapy (HIPEC) protocols lacks effectiveness *in vitro* and *ex vivo*

**DOI:** 10.1101/709055

**Authors:** Markus W. Löffler, Nick Seyfried, Markus Burkard, Benedikt Oswald, Alexander Tolios, Can Yurttas, Franziska Herster, Joseph Kauer, Tarkan Jäger, Karolin Thiel, Sebastian P. Haen, Hans-Georg Rammensee, Sascha Venturelli, Matthias Schwab, Alfred Königsrainer, Stefan Beckert

## Abstract

Cytotoxicity of oxaliplatin-containing solutions (OCS), sampled during patient treatment with hyperthermic intraperitoneal chemotherapy (HIPEC), was assessed by well-established continuous impedance-based real-time cell analysis (RTCA) *ex vivo*. HIPEC treatment was replicated by exposing OAW-42 cancer cells to OCS for 30 or 60 minutes at 42 °C. In contrast to previous observations with continuous exposure, where cytotoxicity was proven, identical OCS obtained during HIPEC did not induce cell death reproducibly and showed strongly attenuated effects after only 30 minutes of application. Based on these unexpected findings, spike-ins of oxaliplatin (OX) into peritoneal dialysis solution (PDS) or dextrose 5 % in water (D5W) were used to replicate HIPEC conditions, as used in either our own protocols or the recently presented randomized controlled PRODIGE 7 trial, where OX HIPEC for 30 minutes failed to produce survival benefits in colorectal carcinoma patients. With OX-spiked into D5W or PDS at identical concentrations as used for PRODIGE 7 or conforming with own HIPEC protocols, we did not observe the expectable cytotoxic effects in RTCA, after replicating OX HIPEC for 30 minutes. These results were corroborated for both solvents at relevant drug concentrations by classical end-point assays for cytotoxicity in two cancer cell lines. Further results suggest that penetration depth, drug dosage, exposure time and drug solvents may constitute critical factors for HIPEC effectiveness. Accordingly, we witnessed substantial cell shrinkage with both PDS and D5W, potentially contributing to reduced drug effects. Based on these results, intensified pharmacological research seems warranted to establish effective HIPEC protocols.

**Key Points:**

- Oxaliplatin (OX)-containing solutions obtained during patient treatment with Hyperthermic intraperitoneal chemotherapy (HIPEC) unexpectedly showed low cytotoxicity in an impedance-based *ex vivo* cytotoxicity cell assay.
- OX cytotoxicity under HIPEC conditions could be enhanced by extending drug exposure to one hour by an impedance-based *ex vivo* cytotoxicity cell assay.
- HIPEC failed to show survival benefits in the randomized controlled PRODIGE 7 trial and was questioned in the aftermath.
- Clinically relevant OX concentrations applied in conjunction with hyperthermia (42 °C) for 30 minutes, as used either at our own medical center or according to the PRODIGE 7 trial, proved predominantly ineffective, when used according to HIPEC routines in an impedance-based *in vitro* cytotoxicity cell assay.
- Respective findings were corroborated in two different cell lines and by two established end-point assays, showing that 50 % cell death could not be reached by the same HIPEC treatment with OX, in contrast to continuous drug exposure.
- As potentially relevant factor, the thickness of the exposed cell layer was identified, requiring at least ~100 µm penetration depth for our model to indicate effectiveness.
- Additionally, we show relevant cell shrinkage by two drug diluents used either at our own medical center or according to the PRODIGE 7 trial, potentially associated with fluid shifts out of the cell and impaired drug effects.
- Our own as well as recent findings by Ubink *et al.* (Br J Surg. 2019. doi: 10.1002/bjs.11206) support the notion that lacking effectiveness of OX HIPEC may explain the negative PRODIGE 7 trial results.

## Introduction

Hyperthermic intraperitoneal chemotherapy (HIPEC) was introduced about four decades ago (1, 2) and it is meanwhile an established treatment option for peritoneal metastasis (PM), in combination with cytoreductive surgery (CRS). This approach of complete macroscopic surgical tumor removal combined with immediate intraoperative heated abdominal chemotherapy lavage was adopted worldwide and is currently most common in the treatment of PM from colorectal (CRC) and ovarian cancers (OvCa) (3). In contrast to CRS, where attempts of standardization do exist (4) and a learning curve for surgical improvement is well-established (5, 6), HIPEC treatment remains very heterogeneous (3, 7). A theoretical basis providing a rationale for adding HIPEC to CRS is the pharmacokinetic advantage of exposing malignant cells directly to high drug concentrations (8) as well as a compartmental effect called *peritoneal-plasma barrier* (9), which presumes that peritoneal malignancies are only insufficiently reached by intravenous (i.v.) chemotherapies.

Historically, PM has been regarded as a palliative disease stage with very limited survival prospects. The first randomized controlled clinical trial (RCT) with CRS + HIPEC vs. systemic chemotherapy showed a significantly improved median overall survival (OS) that increased from 12.6 months to 22.3 months in the CRS + HIPEC group in CRC (10). In this trial by Verwaal *et al.*, HIPEC treatment involved mitomycin c (MMC) exposure for 90 minutes and the treatment was associated with long-term benefit for some selected patients (11). Among the most frequently used drugs for PM in CRC is further oxaliplatin (OX) (12). Since CRS + HIPEC constitute a complex multifactorial compound treatment, the contribution of each of the components was unclear so far, in particular whether there is any added value by HIPEC treatment. A current RCT in PM from CRC answered this question for OX HIPEC over 30 minutes, showing an impressive unexpected median OS of ~41 months for CRS but failed to reach any survival benefit by adding HIPEC (13). In this trial by Quenet *et al.*, HIPEC dosage was standardized to 460 mg OX per every two litres of solvent (i.e. 230 µg/ml ≙ 579 µM) with open and 360 mg (i.e. 180 µg/ml ≙ 453 µM) with closed HIPEC (14). Based on recent positive findings in a RCT by van Driel *et al.* in OvCa patients, where an OS benefit could be established for 90 minutes cisplatin-based HIPEC (15), the treatment rationale seems to remain valid for selected patients. On this background, the pressing question arises, why OX HIPEC has performed relatively poor? Of note, there are fundamental differences in study designs and the implications of different HIPEC protocols are poorly understood so far. Consequently, all women included in the van Driel *et al.* trial received cisplatin HIPEC and had responded to neo-adjuvant platinum-based chemotherapy, in addition to an extended drug exposure time compared to the PRODIGE 7 trial.

Since OX containing solutions (OCS) obtained during our own HIPEC procedures did not induce expectable cell death and in order to investigate the negative results obtained in the PRODIGE 7 trial (13) on a pharmacodynamic level, we adopted *inter alia* a previously established continuous real-time cell assay (16) to evaluate OX-based HIPEC treatment in a model system. Supporting our findings concerning the lacking effectiveness of short-term drug exposure, a recent study modelling HIPEC in CRC-derived organoids confirmed generally weak drug effects, failing to reach 50 % tumor cell elimination after 30 minutes OX exposure (17).

Here, we show additional experimental data in a model, indicating that the OX-based HIPEC protocol applied in the PRODIGE 7 trial as well as according to our own protocol may fail to achieve an effective elimination of residual malignant cells within the peritoneum. Further, we evaluate some potential causes for the observed ineffectiveness of HIPEC treatment with OX.

## Results

### Exposure to intraoperatively sampled oxaliplatin-containing solutions (OCS) for 30 minutes frequently fails to induce 50 % cell death (LC_50_) in an *in vitro* model system

We used a continuous label-free impedance-based real time cell analysis (RTCA) system to assess the effects of applying OCS obtained during HIPEC to ovarian cystadenocarcinoma (OAW-42) cells, as previously established (16). We have shown that this method enables monitoring cell death with a good temporal resolution, following up on cell viability after treatment. However, this time we intended to precisely model HIPEC conditions and used OCS sampled during HIPEC, exposing cells for 30 and 60 minutes at 42 °C, subsequently washing with phosphate-buffered saline (PBS) and exchanging OCS for serum supplemented cell culture medium.

Since we had previously tested respective OCS obtained from patients sampled during HIPEC treatment (16), we were interested in how this well-characterized sample material would perform in an *ex vivo* RTCA assay, recreating the precise conditions prevailing during HIPEC. We have shown prior that OCS obtained from HIPEC in nine patients invariably induced complete cell death, when OAW-42 cells were incubated with OCS supplemented to cell culture medium within three days (16). Of note, in those assays OCS were diluted by 50 % with serum complemented medium to enable sustaining the cell culture. Due to this technical requirement, resulting drug concentrations were bisected as compared to OX amounts encountered during HIPEC *in vivo*. Thus, based on the drug concentrations administered (on average 90 µg/ml ≙ 227 µM) for HIPEC, we can assume respective OX concentrations did not exceed about 45 µg/ml (≙ 113 µM) in any of the experiments. Nevertheless, resulting OX concentrations sufficed to eliminate exposed tumor cells, results that were corroborated by additional OX spike-in experiments performed in parallel (16). Moreover, it has been established that only 0.05 µM OX suffice to kill 50 % of exposed OAW-42 cells (LC_50_) within 72 hours (18).

In our hands however, the identical OCS obtained during HIPEC in those nine patients and analyzed previously (16) were unable to consistently induce cell death in our *ex vivo* model system, when exposing cells to OCS for 30 minutes at 42 °C. Strikingly, in 6/9 investigated patients there was no discernible induction of cell death and the treatment proved ineffective for reaching 50 % reduction of impedance (LC_50_) within three days (Fig. 1a). A prolongation of exposure time to one hour could not rescue the effects in all patients but it strongly improved LC_50_ attainment, favouring early time points after HIPEC initiation (Fig. 1b).

**Figure 1.**
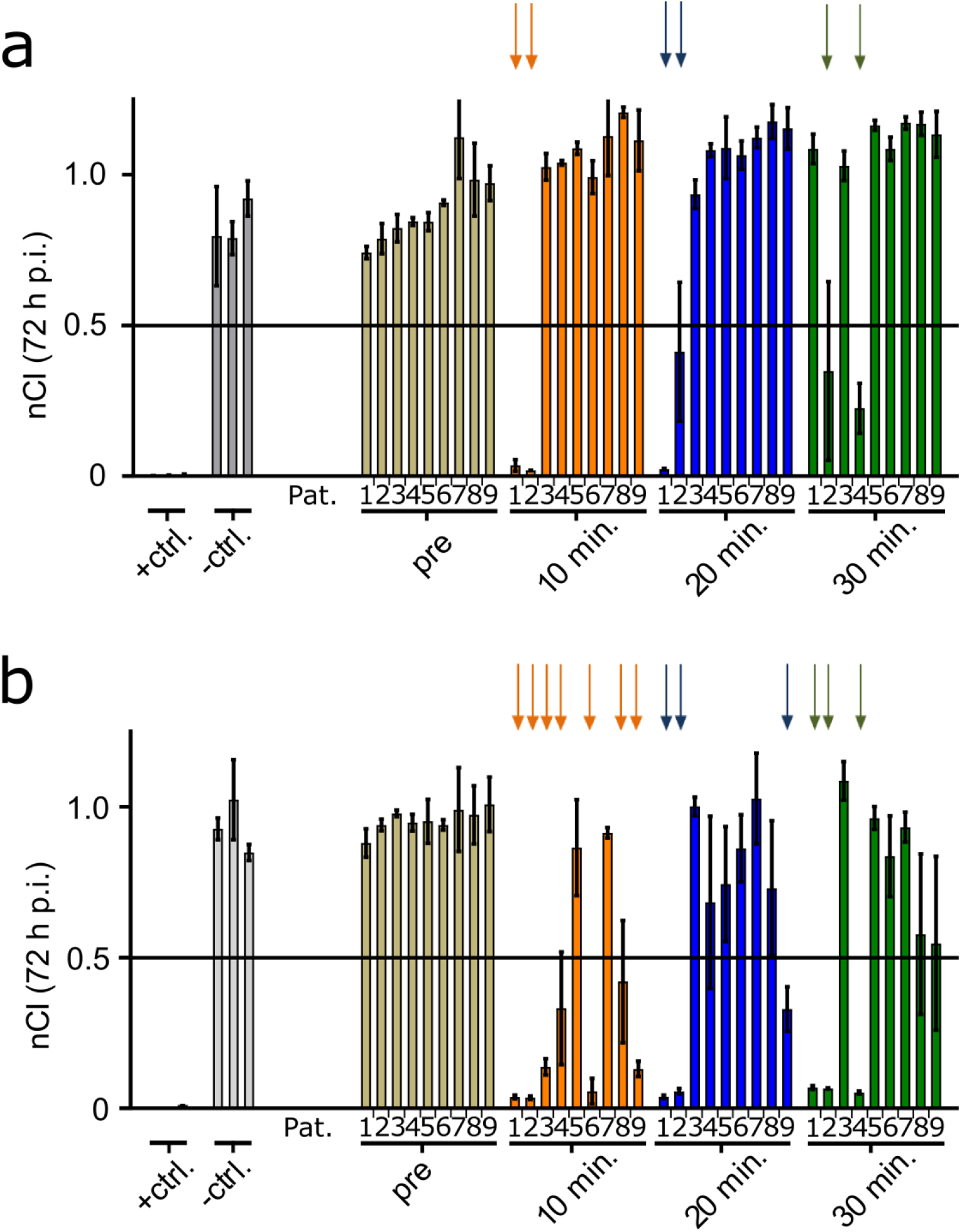
Cell exposure for 30/ 60 minutes at 42 °C in a time resolved in vitro model system using oxaliplatin-containing solutions (OCS) obtained from patients during HIPEC, assessing drug effects at three days after treatment by impedance based real time cell analysis (RTCA). Normalized cell index (nCI; normalized to 1 at treatment) from RTCA measurements at three days (72 hours) after short-term incubation with hyperthermia and OCS (*post interventionem*; p.i.) obtained from patients during HIPEC. From left to right: (+ ctrl.): Triton; (-ctrl.): Physioneal 40; pre (circulated solvent [Physioneal] without addition of OX), and OCS sampled 10, 20 and 30 minutes after addition of OX to the circuit during HIPEC treatment. Pat. 1-9 are depicted from left to right as already characterized in (16). At HIPEC initiation OCS samples contained a theoretical average OX dosage of 90 µg/ml (≙ 227 µM). Panel **(a)** shows effects after 30 minutes incubation at 42 °C with respective OCS. Panel **(b)** shows respective effects after 60 minutes OCS incubation at 42 °C. Colored arrows mark conditions, when an average decrease in impedance of >50 % of the initial values (LC_50_) was reached 72 hours after treatment. Graphs show mean ± SD (of 2-5 replicates). The LC_50_ threshold is marked with a black line. A decrease of nCI values signifies cell death of OAW-42 target cells. Respective normalized cell index (nCI) readings in 6-hour intervals of RTCA measurements are provided in Fig. 2 as well as **Suppl. Fig. 1-14**.

**Figure 2.**
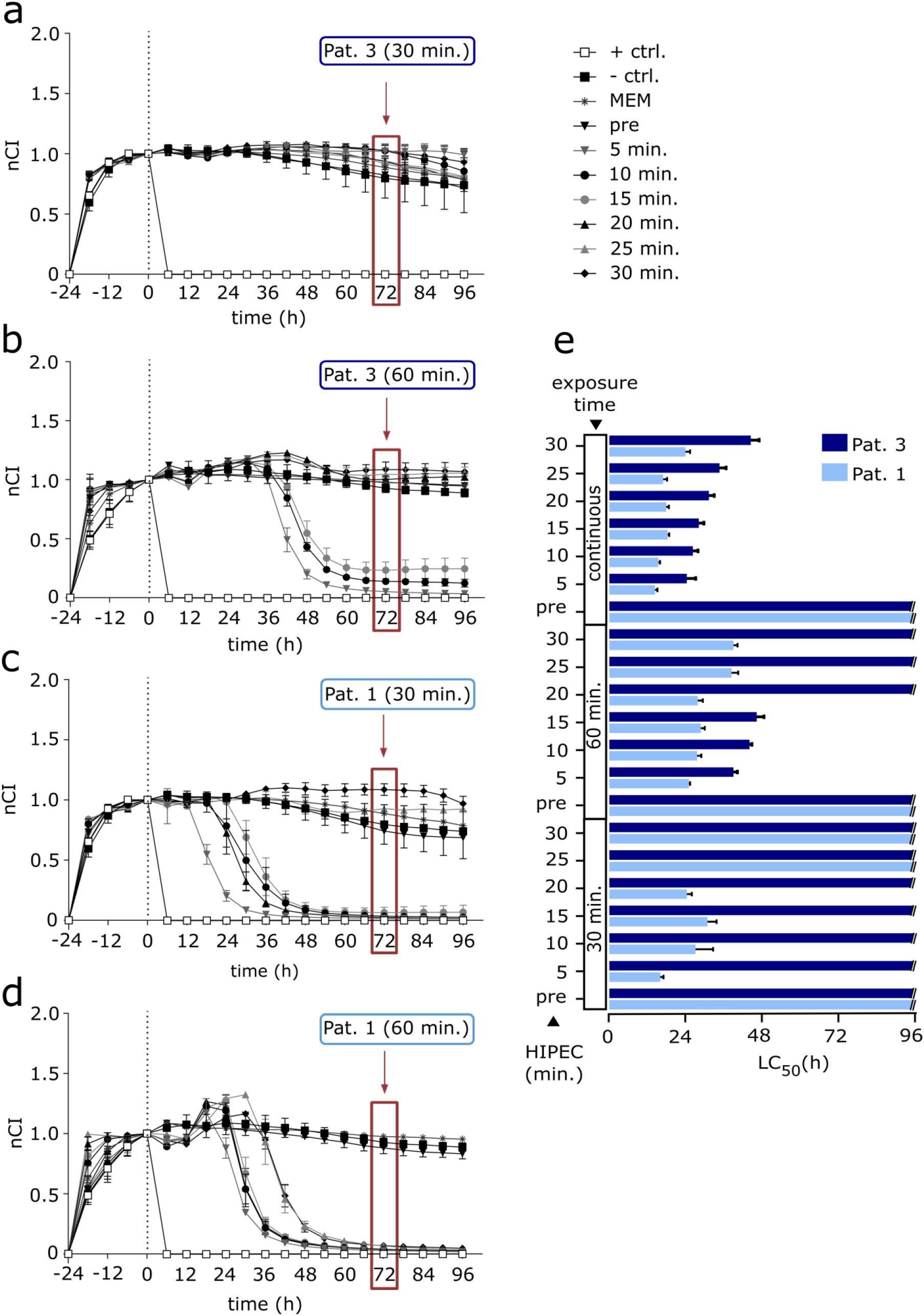
Cell exposure for 30 or 60 minutes at 42 °C in a time resolved in vitro model system using oxaliplatin-containing solutions (OCS) obtained from patients during HIPEC. Normalized cell index (nCI) in 6-hour intervals from RTCA measurements of platinum-sensitive OAW-42 cells, incubated with OCS (0 h) at 42 °C, previously obtained from patients during HIPEC for the indicated periods and drug solvent circulated through the abdomen sampled before drug application (pre). OCS was obtained at time points 5, 10, 15, 20, 25 and 30 minutes (min.) after drug addition to the HIPEC circuit. Controls represent: (+ ctrl.): Triton; (-ctrl.): Physioneal 40 or medium (MEM). Impedance values (nCI) of samples obtained during HIPEC from Pat. 3 incubated for 30 minutes **(a)** and 60 minutes **(b)**, as well as from Pat. 1 after incubation for 30 minutes **(c)** and 60 minutes **(d)** are shown. The duration until 50 % impedance decrease (LC_50_) was reached with 30/ 60 minutes exposure or continuous incubation of respective samples is given **(e)**. The samples from patients (pre + OCS) correspond to the material previously characterized and values of continuous incubation were taken from Löffler *et al.* (16). This material contains a theoretical maximum average OX dosage of 90 µg/ml (≙ 227 µM), however for continuous exposure samples had to be diluted by 50 % with serum supplemented medium due to technical reasons. Values were normalized to 1 at the start of treatment (0 h). The red box highlights values three days (72 hours) after treatment (see also Fig. 1). A decrease of nCI values signifies cell death of OAW-42 target cells. Graphs show mean ± SD (of 2-4 replicates). Respective normalized cell index (nCI) readings in 6-hour intervals of RTCA measurements after incubation with HIPEC sample materials obtained from Pat. 2 and 4-9 are provided in **Suppl. Fig. 1-14**.

Indeed, there were striking differences between different samples (i.e. OCS ≙ Physioneal peritoneal dialysis solution containing OX at 300 mg/m² body surface area; resulting in a calculated mean concentration of 93.7 µg/ml, ranging from 75.0–108.9 µg/ml (16)) obtained during HIPEC. Some OCS showed no cytotoxic effects in OAW-42 cells after 30 minutes exposure at 42 °C (Fig. 2a), which could be rescued by prolonged exposure in some cases (Fig. 2b). OCS from different patients showed effects already after 30 minutes exposure instead (Fig. 2c). Again, in this latter case cytotoxic effects were boosted by longer exposure to OCS (Fig. 2d). A comparison with published results of continuous OCS exposure, inducing complete cell death and reaching LC_50_ unanimously within 72 hours, even when reducing the drug concentrations by 50 % (16), emphasizes treatment related differences. Samples taken earlier during HIPEC showed a tendency to stronger effects and longer exposure proved more effective for reaching LC_50_ values (Fig. 2e), whereas several OCS fail to induce any discernible cell death after 30 minutes of exposure. A complete dataset is provided in **Suppl. Fig. 1-14**.

### Oxaliplatin (OX) exposure for 30 minutes fails to induce cell death in a time resolved *in vitro* model system

To recreate the conditions prevailing during HIPEC as authentically as possible, we exposed OAW-42 cells to defined OCS, spiked with OX in a dose range from 5.6 – 230 µg/ml (≙ 14 µM – 579 µM), encompassing the exact concentrations calculated from reports of the PRODIGE 7 trial (180 µg/ml ≙ 453 µM for closed HIPEC and 230 µg/ml for the open (coliseum) procedure; (13, 14)) as well as based on data from own HIPEC conduct (~90 µg/ml ≙ 227 µM) (16). This dose range is also in line with previous published HIPEC *in vitro* models (17).

In order to recreate the HIPEC setting, cells were exposed to OCS for 30 minutes at 42 °C, either spiked into peritoneal dialysis solution (PDS; Physioneal 40) as used in procedures at our Tübingen University Hospital or into dextrose 5 % in water (D5W) as used in the PRODIGE 7 trial. Subsequently, drugs were removed, cells were washed and cultured in serum supplemented medium again.

Of note, we used a dense cell layer in all experiments and a positive readout required drug penetration into deeper cell layers. A titration series revealed a linear relationship between the seeded cell numbers and the resulting cell layer thickness, showing an average diameter of 94 ± 5 µm (**Suppl. Fig. 15**) for 50.0 × 10^3^ cells seeded per well in a 96-well format.

Using this cell culture model, we did not observe any treatment induced cell death induced when diluting OX in D5W up to a concentration of 230 µg/ml during three days of follow-up (Fig. 3a), whereas the positive control (1 % Triton X-100) proved instantaneously effective for a complete destruction of the cell layer. It should be pointed out that this method is limited to measuring a complete destruction of the full thickness cell layer and partial effects may not result in any measurable results, as the electrodes for detection are located at the well bottom. Since at our hospital and other medical centres OX is diluted in PDS (7, 16), such as Physioneal 40 (containing 2.27 % dextrose), we additionally tested OAW-42 cells under identical experimental conditions with OX diluted in PDS. Using PDS as a diluent, we evidenced that the maximum concentration of 230 µg/ml OX (≙ 579 µM; corresponding to the dosage used in the PRODIGE 7 trial) was effective in inducing cell death (Fig. 3b), whereas effects stayed far behind of continuous exposure of OX spiked into medium (Fig. 3c). Continuous exposure showed a dose dependency and LC_50_ was reached at much lower concentrations than evidenced with short term exposure. OX exposure prolonged to 60 minutes showed to be effective only for 230 and 180 µg/ml when spiked into PDS (**Suppl. Fig. 16**), whereas 60 minutes exposure with OX spiked into D5W proved technically unfeasible due to cell detachment, potentially due to osmotic effects caused by D5W.

**Figure 3.**
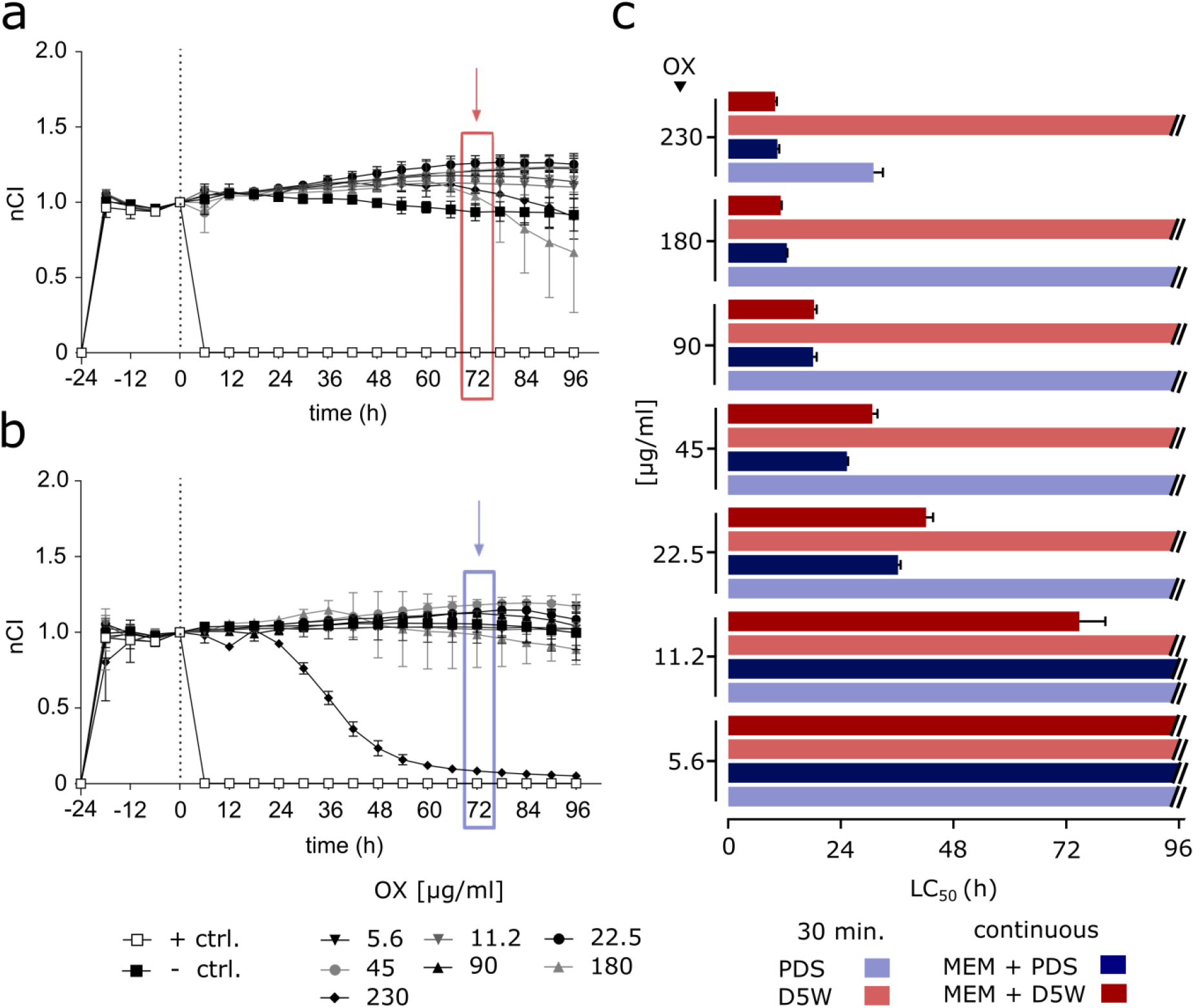
Cell exposure to oxaliplatin at 42 °C for 30 minutes in a time resolved in vitro model system. Normalized cell index (nCI) in 6-hour intervals obtained from RTCA impedance measurements of platinum-sensitive OAW-42 cells incubated for 30 minutes at 42 °C with the specified amounts of OX spiked into dextrose 5 % in water (D5W) **(a)** or Physioneal 40 (PDS) **(b)** performed at time point 0 hours (h). (+ ctrl.): Triton; (-ctrl.): Physioneal 40; OX (oxaliplatin); PDS (peritoneal dialysis solution; Physioneal 40); RTCA (real-time cell analysis). Graphs show mean ± SD (of 2-3 replicates). The duration until 50 % impedance decrease (LC_50_) was reached after 30 minutes exposure of OX spiked into PDS/ D5W or with continuous incubation of OX spiked into D5W or PDS (together with 50 % serum supplemented medium) at the given concentrations is depicted **(e)**. Values were normalized to 1 at the start of treatment (0 h). The colored boxes highlight values three days (72 hours) after treatment. Respective nCI in 6-hour intervals obtained from RTCA impedance measurements of platinum-sensitive OAW-42 cells incubated continuously with OX are provided as **Suppl. Fig. 17 & 18**.

### Oxaliplatin (OX) exposure for 30 minutes frequently fails to induce 50 % cell death (LC_50_) in end point assays

To verify these unexpected findings by independent experimentation and in different experimental assays, we used both a robust fluorometric assay (CTB; CellTiter-Blue^®^) based on the conversion of resazurin to resorufin, occurring only in living cells (19), and sulforhodamine B (SRB), which binds stoichiometrically to proteins under mild acidic conditions, constituting an assay well-established for investigating cytotoxicity (20, 21). To recreate the conditions of the PRODIGE 7 trial (13), we first spiked OX into a D5W solution, exposing cells to OCS for 30 minutes at 42 °C and subsequently cultured the washed cells in medium for 72 hours before analysis. Cells were intentionally seeded densely to reach comparable conditions as used previously in the RTCA assays. To broaden our analysis we also added the human colon carcinoma cell line HT29 (22), which has been included in the *NCI60 human tumour cell line anticancer drug screen* panel and is therefore been very well characterized (23). The LC_50_ for OX in HT29 has been established at 72.44 µM (24).

In the CTB assay both the OAW-42 and the HT29 cell lines showed no reduction of mean cell viability below 50 % (LC_50_) three days after treatment, in none of the assessed conditions (Fig. 4a). The robustness of these findings was confirmed by reproducing respective results with the SRB assay. Repeating experiments under identical conditions with OX spiked into PDS failed again to induce LC_50_ at three days following treatment for all tested conditions (Fig. 4b). This was also evidenced for OX spiked into Ringer’s lactate solution (RL) (**Suppl. Fig. 19**), even though drug effects seemed slightly more pronounced than those observed with OX spiked into D5W and PDS. In contrast, when OX was spiked into culture medium (to allow for extended cultivation) and remained with the cells, both cell lines (OAW-42 and HT29) showed clearly enhanced induction of cell death. Cell viability reproducibly decreased to <50 % (LC_50_) within three days, when exposed to drug concentrations relevant for HIPEC (≥ 90 µg/ml ≙ 227 µM) with both assays and cell lines (Fig. 4c).

**Figure 4.**
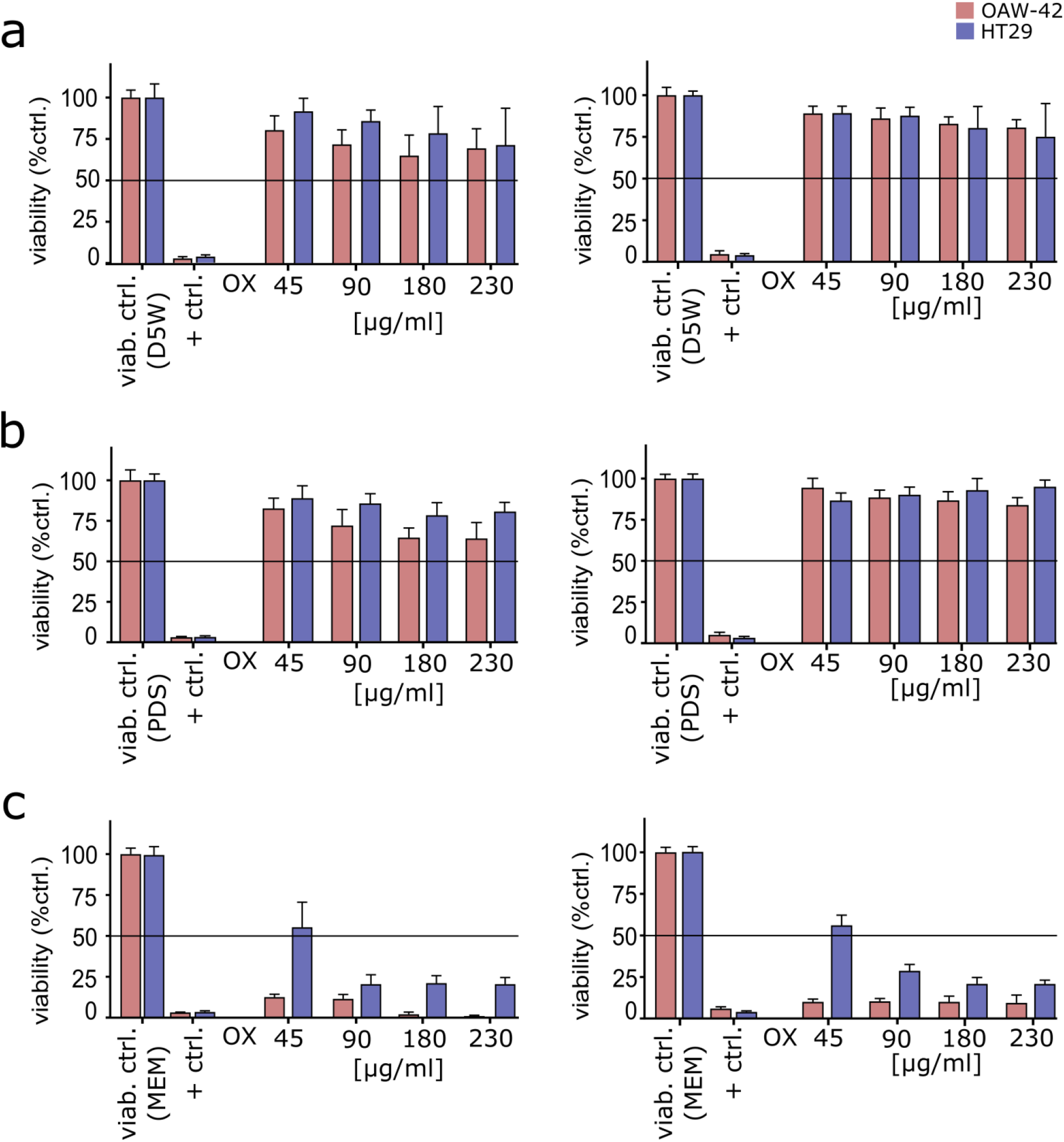
Cell death after incubation of OAW-42 and HT29 cells with OX spiked into dextrose 5 % in water (D5W) or peritoneal dialysis solution (PDS) for 30 minutes at 42 °C and continuous exposure measured by end point assays. Platinum-sensitive OAW-42 cells and the colon carcinoma cell line HT29 were seeded at a density of 3.15 × 10^5^ cells per well (in a 24-well plate) and incubated for 30 minutes at 42 °C with the specified amounts of OX spiked either into dextrose 5 % in water (D5W) **(a)**, or into PDS **(b)**. Further, cells were incubated continuously with the specified amounts of OX spiked into culture medium (MEM) to allow for continuous drug exposure and heated likewise initially (i.e. 30 minutes incubation at 42 °C followed by 72 hours at 37 °C) **(c)**. After OX exposure, cells were washed and subsequently cultured in serum supplemented medium for another three days or remained with drugs in culture medium (MEM). Afterwards, a fluorometric resazurin-based (CellTiter-Blue^®^, CTB) (**left graph**) and a sulforhodamine B (SRB) assay (**right graph**) was used to determine cell viability. Cells were normalized to cells treated identically with D5W/ PDS/ MEM only (viability ctrl.). (+ ctrl.): Triton. Graphs show mean ± SD from three independent experiments each performed in triplicates. The LC_50_ threshold is marked with a black line.

### Short-term exposure to drug diluents used in HIPEC for 30/ 60 minutes causes significant cell shrinkage in OAW-42 cells

Although high sugar containing aqueous solutions such as D5W and PDS are often used for OX dilution in HIPEC (7), this practice has been connected with frequent adverse events including post-operative hyperglycemia and/ or electrolyte disturbances (25) and was questioned based on pharmacological grounds (26). Nevertheless, since both D5W and PDS are hypertonic solutions, we assumed they may affect directly exposed OAW-42 cells based on their physiological properties. To this end, we exposed cells to D5W and PDS under conditions comparable to HIPEC (42 °C) and measured cellular appearance by flow cytometry. We found that 30 minutes exposure suffices to cause significant cell shrinkage with both D5W and PDS, compared to untreated cells (Fig. 5a), which is further enhanced after 60 minutes (Fig. 5b). This substantial decrease in cell size as proven by flow cytometry may be explainable by a relevant fluid shift to the extracellular space.

**Figure 5.**
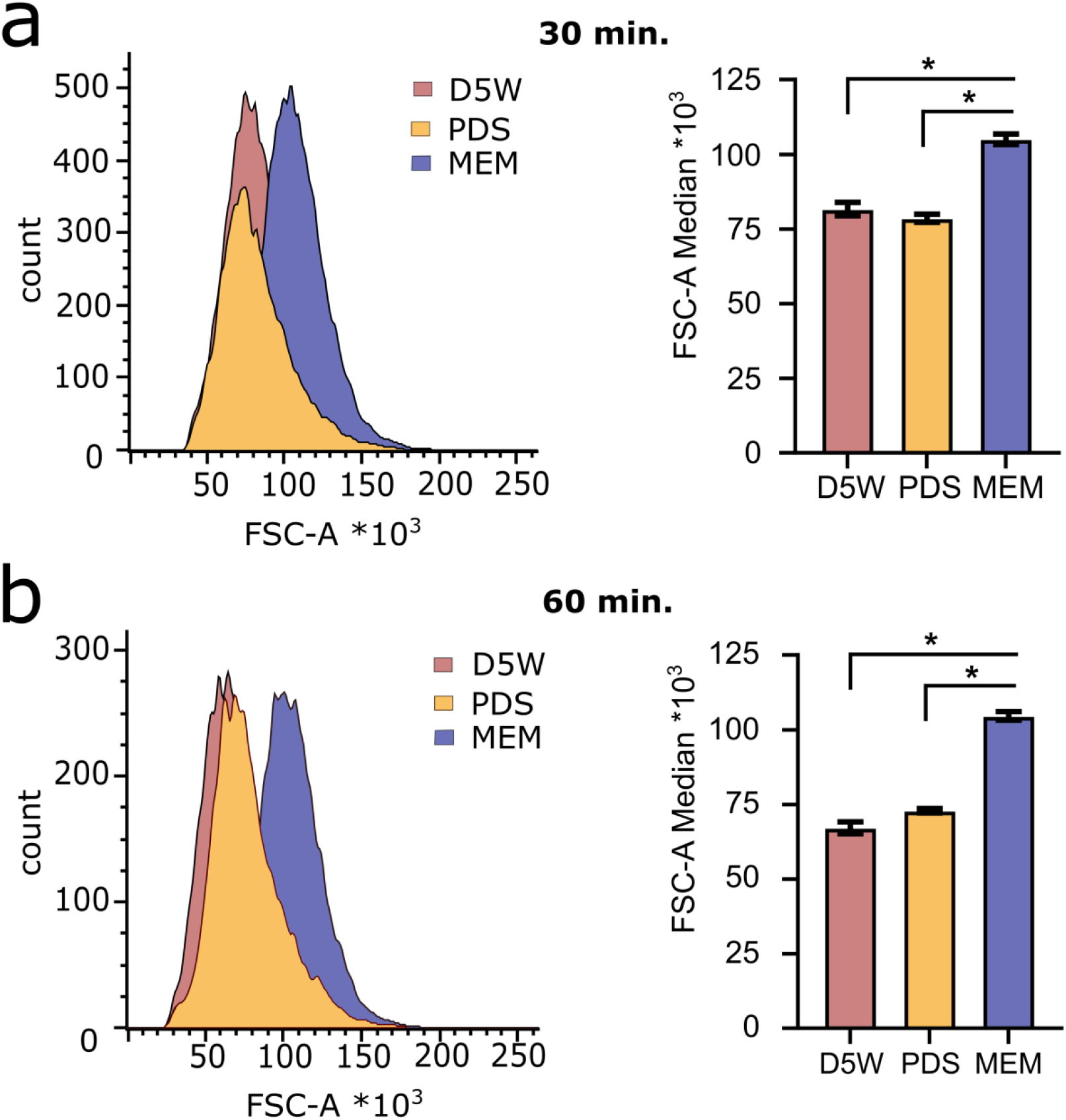
Exposure of OAW-42 cells to D5W or PDS for 30 minutes at 42 °C leads to relevant cell shrinkage and is enhanced after 60 minutes. Platinum-sensitive OAW-42 cells were incubated with 5 % dextrose in water (D5W) or with peritoneal dialysis solution (PDS) at 42 °C for 30 minutes **(a)** or 60 minutes **(b)** and compared to untreated cells cultured in medium (MEM). Histograms from flow cytometry showing cell counts vs. forward scatter area (FSC-A) (**left panel**) and a comparison of FSC-A between the different conditions (**right panel**). Significant differences are marked by an asterisk (Bonferroni corrected Student’s *t*-test).

## Discussion

Previously, we have shown that the OCS used for HIPEC, which have been obtained from patients during 30 minutes of treatment, retain their cytotoxic properties (16). In respective experiments, all tested OCS samples induced complete cell death *ex vivo* in a relatively thick cell layer (containing 50.0 × 10^3^ cells/ well; ~100 µm thickness) after OCS exposure, unanimously reaching >50 % cell death (LC_50_) within three days. Due to technical reasons, only half the concentrations used for HIPEC *in vivo* could be applied for *ex vivo* RTCA assays in these experiments, enabling the addition of serum supplemented medium to sustain cell culture. Results showed that cytotoxic properties of HIPEC solutions were only slightly attenuated in those samples previously circulated through patients’ peritoneal cavities for longer. Intriguingly, even samples spiked with about ^1^/_10_ of the highest OX concentration that was used in the PRODIGE 7 trial (13) were able to eliminate the exposed full thickness cell layer. Of note, when characterizing these OCS obtained during HIPEC from nine patients (theoretically containing a maximal average OX amount of 90 µg/ml ≙ 227 µM), a relatively steady fraction of roughly 50 % of the initially administered OX was detectable by our mass spectrometry method (16). Measurements yielded a compound most probably formed by the reaction of OX with sodium bicarbonate contained in the PDS used as a vehicle for abdominal perfusion. This compound further constituted the predominant fraction of platinum compounds detectable by this method, supporting the notion that the drug solvent may be a relevant influencing factor in HIPEC.

Re-testing these well-characterized patient samples (16), we aimed to precisely model HIPEC conditions encountered *in vivo*. To our knowledge, a comparable model has not been established before and mainly animal models have been used for pre-clinical characterization of HIPEC effects (27, 28), whereas realistic *in vitro* and *ex vivo* models were lacking. Hence, here we aimed at recreating precise HIPEC conditions. To this end, we exposed a 100 µm thick cell layer (corresponding to ~ ⅓ of the diameter of a grain of salt) to OCS for 30 minutes at 42 °C, subsequently washing cells and incubating them in fresh serum containing medium. Surprisingly, only OCS from 3/9 patients showed marked cell death under these conditions and all the OCS samples obtained from six patients did not even reach LC_50_ values (50 % decrease of cell viability) within 72 hours, when measured at the bottom cell layer. Further, drug effects varied strongly in samples taken at different time points. Nevertheless, when using the identical sample material drug efficacy could be relevantly boosted by one-hour OCS incubation.

In the context of the recently presented randomized controlled PRODIGE 7 trial (13), which tested CRS only vs. CRS and HIPEC (OX for 30 minutes) in PM from CRC, failing to show any difference in overall survival (OS) during five years of follow-up, we were very interested in also modelling respective treatments in our HIPEC model. However, the PRODIGE 7 trial used both 2.0-2.5-fold higher OX concentrations than those employed at our medical center and a different solvent, namely 5 % dextrose in water (D5W), yielding final OX concentrations at 180 – 230 µg/ml (≙ 453 – 579 µM). Still, both HIPEC treatment schedules are well-comparable (OX exposure for 30 minutes and hyperthermia). We therefore aimed to simulate the PRODIGE 7 treatment conditions and spiked OX into D5W over a wide concentration range. Further, we spiked OX into PDS (Physioneal 40), which was used at our medical center for HIPEC, encompassing the concentrations used for the PRODIGE 7 RCT and in our patients. Interestingly, all tested conditions proved insufficient to induce deep penetrating cell death after 30 minutes exposure, except for the highest concentration of OX when combined with PDS. Two classical endpoint assays (CTB and SRB (19–21)) generally verified these findings in OAW-42 cells as well as in colon cancer cells (HT29). Overall, in our assays both RTCA and endpoint assays (CTB/ SRB) showed strongly attenuated effects of short-term exposure to OX spiked into either D5W or PDS (as well as Ringer’s lactate), whereas continuous exposure to OX generally reached LC_50_ values in all conditions tested. Additional evidence suggesting that short-term OX use is insufficient to eliminate small CRC metastases, has recently been provided in patient-derived organoids from five patients, where 50 % cell death was never reached after modelling HIPEC over a wide clinically relevant concentration range (17).

Further, significant cell shrinkage was observed in cell culture after short-term exposure to both D5W and PDS, theoretically expectable from hypertonic solutions, likely caused by a fluid shift into the extracellular space. This aspect, although not sufficiently investigated yet, may contribute to explain our findings, together with a very limited penetration depth of OX, complex pharmacology (29) and short-term exposure.

It should be emphasized that the PRODIGE 7 trial has excelled with a median OS of ~41 months in both treatment groups (13), exceeding the anticipated OS by far, which proves that CRS is indeed a highly effective treatment option for PM of CRC.

On the downside, this HIPEC protocol showed an inferior economic and adverse events profile, increasing morbidity and hospital stay. If this notion also applies to other drugs used for HIPEC in the same way is unclear to date. Indeed, a recent RCT by van Driel *et al.* in OvCa with cisplatin HIPEC for 90 minutes resulted in superior OS compared to CRS only, validating the general concept, *alas*, in a highly selected group of women with cancers proven to be sensitive to neo-adjuvant platinum-based chemotherapy (15).

From a pharmacological point of view the OX results may not come very surprisingly, since monotherapy showed very limited effectiveness in CRC already, when administered by i.v. route (with response rates between 12 and 24 % (30), rendering the drug primarily a candidate for poly-chemotherapy. To mitigate this, in the PRODIGE 7 trial OX administration to the abdomen was combined with 5-FU and leucovorin i.v., presuming bidirectional drug effects (31). This practice may be disputed, when assuming that the frequently postulated concept of a *peritoneal-plasma barrier* is true (9). Of note, OX is a particularly complex cytotoxic agent with triphasic elimination kinetics in circulation (32) and features a short plasma half-life (33), forming various intermediates *in vivo* (34), some of which still remain uncharacterized. OX stability in aqueous solutions is explained by concentration effects and has been determined at no less than three years in D5W, forming association complexes at high concentrations that may affect uptake and molecular drug mechanisms (35). Its physicochemical stability in combination with D5W at concentrations >200 µg/ml (≙ ~500 µM) may further contribute to chemical inertness (36).

In this context, we might also challenge the notion of OX inefficiency in chloride containing solutions (16, 26), as these form (instable) intermediates with OX but retain their required cytotoxic properties (37). OX derived monochloro- and dichloro-compounds have even been suggested to exert enhanced cytotoxicity (26). Dextrose-containing HIPEC vehicles instead may cause hyperglycemia (38) and electrolyte disturbances, which may influence postoperative morbidity and mortality (26) and have effects such as fluid shifts that may thwart the treatment.

Regardless, we assume the most important factor to be considered is drug penetration into cell layers, which is the best explanation for the RTCA findings and the enhanced drug effects after prolonged incubation even at substantially lower dosage. It should be considered that OX cytotoxicity is well-established for the used cell lines at a small fraction of the concentrations employed here (18, 24). Since we used cell culture without substantial intercellular matrix or surrounding tissue, this may even become more relevant *in vivo* as already supposed in the past (8).

## Conclusion

Our results and previous work underscore the need for basic pharmacological research in HIPEC enabling to discern relevant factors in a complex multifaceted treatment. These findings suggest that drug distribution into deeper tissue layers may be a crucial factor as well as exposure time and dosage but also the vehicle used for OX dilution in HIPEC may relevantly alter drug induced effects.

The presented work may not only support the notion that OX has been used too short for HIPEC but instead is a very complex drug that has serious limitations for intraperitoneal application. Based on the restricted penetration depths of most HIPEC drugs (8), this calls for intensified preclinical research and underscores that HIPEC – in contrast to CRS – remains an experimental treatment for peritoneal metastasis of CRC urgently requiring further investigations in well-designed clinical trials as well as basic research. Moreover, it seems imperative to develop or select suitable drugs for the HIPEC setting that are adequate for intraperitoneal application.

## Materials and Methods

### Patient materials

Studies on patient materials were approved by the institutional review board at Tübingen University. All patients gave their written informed consent before study inclusion. Patient characteristics can be assessed elsewhere (16). Drug solvent circulated through the abdomen of patients was collected prior to addition of OX (0 min) and 5, 10, 15, 20, 25, and 30 minutes after starting the HIPEC procedure with OX. For Pat. 8 and 9, samples were only obtained in 10-minute intervals. Samples were stored at −80 °C until usage. Cellular debris and other impurities were cleared by centrifugation (five minutes at 13,000 x g) before analysis.

### Impedance-based real-time cell analysis (RTCA)

Materials and methods were used as established before (16), with slight modifications.

As described previously, platinum-sensitive OAW-42 cells (European Collection of Authenticated Cell Cultures, Salisbury, UK) were grown under appropriate conditions, seeded and used as a model system to monitor cytotoxicity of OX in real time (16). In brief, cells were grown in culture medium (Dulbecco’s Modified Eagle Medium [DMEM], high glucose; Gibco/Life Technologies, Carlsbad, CA, USA, adding 10 % fetal calf serum (FCS); Sigma-Aldrich/Merck Life Science, St. Louis, MO, USA; 100 U/ml penicillin G; PAA, Pasching, Austria; and 100 µg/ml streptomycin; PAA). Cell cultures were periodically tested for mycoplasma using commercially available polymerase chain reaction (PCR) kits (Minerva Biolabs, Berlin, Germany).

A RTCA device (xCELLigence SP; Roche, Grenzach-Wyhlen, Germany) was used as previously described (16). To model HIPEC conditions, after calibration of the device with 100 µl cell culture medium (blank values), 50.0 × 10^3^ OAW-42 cells/ well were seeded in dedicated 96-well plates (E-plate 96; ACEA, San Diego, CA, USA) and left to adhere for 24 hours. Subsequently, fluid was discarded from wells and adherent cells were washed once with PBS (DPBS; Gibco Life Technologies, Carlsbad, CA, USA). Then, 200 µl of peritoneal perfusate obtained during HIPEC from patients as well as defined concentrations of OX (Oxaliplatin-GRY/ oxaliplatin 5 mg/ml; Teva, Petach Tikwa, Israel/ Fresenius Kabi, Bad Homburg, Germany) ranging from 5.6 µg/ml to 230 µg/ml (≙ 14 – 579 µM) spiked into peritoneal dialysis fluid (PDS; Physioneal 40 Glucose 2.27 % m/v; Baxter, Deerfield, IL, USA) or dextrose 5 % in water (D5W; Glucosteril 5 %; Fresenius Kabi) were added. Likewise, adequate controls were employed, including 1 % (v/v) Triton X-100 (Sigma) as a positive (dead) control and negative controls with either PDS or culture medium. HIPEC conditions were subsequently simulated by incubation of cells for 30 or 60 minutes at 42 °C in an ambient air incubation shaker (Infors, Bottmingen, Switzerland) with slight movement (50 rotations per minute). Afterwards, liquids were discarded by flicking and cells were washed twice in PBS and then cultivated in culture medium supplemented with FCS. All samples were analyzed at least in duplicates and impedance was measured continuously in 15 minutes intervals. Technical errors and outliers were removed.

For inter-experiment comparability, the cell index was set to 1 at addition of compounds/ treatment (normalized cell index [nCI]). Results were analyzed using RTCA software (V. 1.2.1). GraphPad prism software (V. 7.01; GraphPad software Inc., La Jolla, CA, USA) was used for presentation of results. Findings of different experiments were combined.

### CellTiter-Blue (CTB) Cell Viability Assay

To estimate effects on cell viability, CellTiter-Blue^®^ Cell Viability Assay (CTB; Promega, Mannheim, Germany) was performed. OAW-42 and HT29 cells were seeded in 24-well plates (3.15 × 10^5^ cells) in a volume of 500 µl as triplicates and incubated overnight at 37 °C and 5 % CO_2_ in a humidified atmosphere. To mimic different HIPEC treatments, medium was discarded, cells were washed with PBS and then incubated for 30 minutes at 42 °C under ambient air conditions with OX (at concentrations of 45, 90, 180 or 230 µg/ml; ≙ 113, 227, 453, 579 µM) diluted either in D5W, PDS, Ringer’s-lactate solution (RL) or cell culture medium. Following treatment, solutions were discarded and replaced by 500 µl of fresh cell culture medium after washing with PBS and incubated (37 °C; 5 % CO_2_) for 72 hours. In parallel, OX-spiked culture medium remained on the cells to verify OX toxicity on OAW-42 and HT29 cells after 72 hours exposure at 37 °C and 5 % CO_2_ in a humidified atmosphere. As controls (+ ctrl.) cell death was induced by lysing cells with 1 % (v/v) Triton X-100 (Roth, Karlsruhe, Germany) for ten minutes immediately prior to CTB staining. To determine cell viability 100 µl of assay reagent was added and gently mixed. Fluorescence measurement was performed one hour after incubation at 37 °C and 5 % CO_2_ for each cell line with the Synergy HT microtiter plate reader (BioTek Instruments Inc., Winooski, VT; record fluorescence (Excitation wavelength: 530/25 and Emission wavelength: 590/35, adjusted sensitivity: 35). CTB assays were repeated in three independent experiments.

### Sulforhodamine B (SRB) Cytotoxicity Assay

OAW-42 and HT29 cells were seeded in 24-well plates (3.15 × 10^5^ cells/ well) in a volume of 500 µl as triplicates and incubated overnight at 37 °C and 5 % CO_2_ in a humidified atmosphere. To mimic HIPEC treatment medium was discarded, cells were washed with PBS and then incubated for 30 minutes at 42 °C under ambient air conditions with OX spiked either into D5W, PDS, RL or culture medium (see concentrations above). After treatment, solutions were discarded and replaced by 500 µl of fresh cell culture medium after washing with PBS and incubated for 72 hours (37 °C and 5 % CO_2_). In a parallel experiment, OX-spiked cell culture medium remained on the cells, to verify OX toxicity on OAW-42 and HT29 cells after 72 hours exposure at 37 °C and 5 % CO_2_. As control (+ ctrl.), cell death was induced by lysing cells with 1 % (v/v) Triton X-100 (Roth, Karlsruhe, Germany) for ten minutes immediately prior to SRB staining. Finally, growth inhibition was evaluated by SRB assay. In brief, medium was discarded, and each well was washed once with ice-cold PBS and fixed with 10 % trichloroacetic acid (TCA) for 30 minutes at 4 °C. After washing with tap water, cells were dried at 40 °C overnight. Then proteins were stained for ten minutes with SRB reagent (0.4 % (w/v) in 1 % (v/v) acetic acid; CAS 3520-42-1, Sigma-Aldrich) and after removing unbound dye with tap water followed by 1 % (v/v) acetic acid, dried again at 40 °C. Protein-bound dye was resolved with 10 mM Tris base (pH 10.5). After ten minutes incubation at room temperature, optical density was measured in triplicates (80 µl volume/ well) in 96-well plates with the Synergy HT microtiter plate reader (BioTek Instruments; measurement wavelength 550 nm, reference wavelength 620 nm). Data represent the mean of optical density values related to medium treated control cells. SRB assays were repeated in three independent experiments.

### Microscopy

In order to obtain a serial dilution, OAW-42 cells were seeded in different densities of 12.5/ 20.0/ 25.0/ 35.0 and 50.0 × 10^3^ cells/ well (96-well plate) and incubated in 200 µl culture medium. After 24 hours of cell culture, the cells were carefully washed with PBS and subsequently fixed with fixation buffer (BioLegend, San Diego, USA) for 10 minutes at room temperature, carefully washed again and stored at 4 °C prior to measurements. Experiments were performed twice.

The thickness of cell layers was obtained measuring z-stacks on a Nikon ti eclipse (by an unbiased observer) using 10x magnification. The analysis was performed with the NIS-Elements (Nikon, Tokyo, Japan) or ImageJ software V. 1.52h.

### Flow Cytometry

OAW-42 cells were seeded in 96-well plates (1 × 10^5^ cells/ well) in either 200 µl of MEM culture medium, PDS, or D5W and kept for 30 or 60 minutes at 42 °C. Immediately prior to flow cytometric analysis, 7-amino-actinomycin D viability staining solution (7-AAD; BioLegend) was added to each well at a final concentration of 600 ng/ml. Dead cells (7-AAD^+^) as well as doublets were excluded. Forward scatter area (FSC-A) served as an indicator for cell size. Samples were analyzed using a FACSCanto II (BD Biosciences, Heidelberg, Germany) and data analysis was performed using FlowJo_V9 software (FlowJo LCC, Ashland, OR). Flow cytometric analysis of cell size under HIPEC conditions was repeated in three independent experiments. Each data point represents the mean value of three replicates.

## Acknowledgments

The authors would like to thank Jan Franko, M.D., Des Moines, Iowa for insights, encouragement and helpful discussions. We also thank Jürgen Winter, Department of General, Visceral and Transplant Surgery, University Hospital of Tübingen, for excellent technical support as well as Jürgen Weinreich, PhD, and Philipp Horvath, M.D., for support with obtaining sample materials.

## Abbreviations

CRC: Colorectal cancer
CRS: Cytoreductive surgery
CTB: CellTiter-Blue® assay
Ctrl.: Control
D5W: Dextrose 5 % in water solution
FSC-A: Forward scatter area
HIPEC: Hyperthermic intraperitoneal chemotherapy
i.v.: Intravenous
LC_50_: 50 % cell viability threshold
MEM: Medium for cell culture
MMC: Mitomycin C
mM: Millimolar
µM: Micromolar
nCI: Normalized cell index
OCS: Oxaliplatin-containing solutions (obtained during HIPEC treatment)
OS: Overall survival
OvCa: Ovarian cancer
OX: Oxaliplatin
PBS: Phosphate-buffered saline
PCR: Polymerase chain reaction
PDS: Peritoneal dialysis solution
PM: Peritoneal metastases
RCT: Randomized controlled trial
RL: Ringer’s lactate
RTCA: Real time cell analysis
SRB: Sulforhodamine B
SD: Standard deviation
TCA: Tichloroacetic acid

