## Supplementary information: Suppl. Figs. 1-19 for "Short-term oxaliplatin exposure according to established hyperthermic intraperitoneal chemotherapy (HIPEC) protocols lacks effectiveness *in vitro* and *ex vivo*"

### Table of contents

**Figure Legends to Suppl. Fig. 1-14** **S3**

**Suppl. Fig. 1** *RTCA Pat. 2: Exposure of OAW-42 cells to OCS for 30 minutes at 42 °C ex vivo* **S4**

**Suppl. Fig. 2** *RTCA Pat. 2: Exposure of OAW-42 cells to OCS for 60 minutes at 42 °C ex vivo* **S4**

**Suppl. Fig. 3** *RTCA Pat. 4: Exposure of OAW-42 cells to OCS for 30 minutes at 42 °C ex vivo* **S5**

**Suppl. Fig. 4** *RTCA Pat. 4: Exposure of OAW-42 cells to OCS for 60 minutes at 42 °C ex vivo* **S5**

**Suppl. Fig. 5** *RTCA Pat. 5: Exposure of OAW-42 cells to OCS for 30 minutes at 42 °C ex vivo* **S6**

**Suppl. Fig. 6** *RTCA Pat. 5: Exposure of OAW-42 cells to OCS for 60 minutes at 42 °C ex vivo* **S6**

**Suppl. Fig. 7** *RTCA Pat. 6: Exposure of OAW-42 cells to OCS for 30 minutes at 42 °C ex vivo* **S7**

**Suppl. Fig. 8** *RTCA Pat. 6: Exposure of OAW-42 cells to OCS for 60 minutes at 42 °C ex vivo* **S7**

**Suppl. Fig. 9** *RTCA Pat. 7: Exposure of OAW-42 cells to OCS for 30 minutes at 42 °C ex vivo* **S8**

**Suppl. Fig. 10** *RTCA Pat. 7: Exposure of OAW-42 cells to OCS for 60 minutes at 42 °C ex vivo* **S8**

**Suppl. Fig. 11** *RTCA Pat. 8: Exposure of OAW-42 cells to OCS for 30 minutes at 42 °C ex vivo* **S9**

**Suppl. Fig. 12** *RTCA Pat. 8: Exposure of OAW-42 cells to OCS for 60 minutes at 42 °C ex vivo* **S9**

**Suppl. Fig. 13** *RTCA Pat. 9: Exposure of OAW-42 cells to OCS for 30 minutes at 42 °C ex vivo* **S10**

**Suppl. Fig. 14** *RTCA Pat. 9: Exposure of OAW-42 cells to OCS for 60 minutes at 42 °C ex vivo* **S10**

**Suppl. Fig. 15** *Assessment of the thickness of an OAW-42 cell layer seeded at different densities* **S11**

**Suppl. Fig. 16** *Oxaliplatin (OX)-spiked into PDS 60 minutes at 42 °C* **S12**

**Supp. Fig. 17** *Continuous exposure of OAW-42 cells to oxaliplatin (OX)-spiked into PDS* **S13**

**Supp. Fig. 18** *Continuous exposure of OAW-42 cells to oxaliplatin (OX)-spiked into D5W* **S14**

**Suppl. Fig. 19** *End point assays: Cell death of OAW-42/HT29 cells exposed to oxaliplatin (OX)-spiked Ringer´s lactate for 30 minutes at 42 °C* **S15**

**Suppl. Fig. 1-14 RTCA Pat. 2 and 4-9:**
**Exposure of OAW-42 cells to OCS for 30/ 60 minutes at 42 °C**

Normalized cell index (nCI) in 6-hour intervals from real-time cell analysis (RTCA) impedance measurements of platinum-sensitive OAW-42 cells, incubated with oxaliplatin-containing solutions (OCS) (0 h) at 42 °C, previously obtained from patients during HIPEC for the indicated periods and drug solvent circulated through the abdomen sampled before drug application (pre). OCS were obtained at time points 5, 10, 15, 20, 25 and 30 minutes after drug addition to the HIPEC circuit (for Pat. 2 and 4-7) and after 10, 20 and 30 minutes (for Pat. 8 and 9). Controls represent: (+ ctrl.): Triton; (- ctrl.): Physioneal 40 and medium (MEM). Impedance values (nCI) of samples obtained during HIPEC from Pat. 2 and 4-9 incubated for 30 minutes (**Suppl. Fig. 1/ 3/ 5/ 7/ 9/ 11/ 13**, respectively) as well as for 60 minutes (**Suppl. Fig. 2/ 4/ 6/ 8/ 10/ 12/ 14**, respectively) are shown below (page S4 - S10). Patient coding corresponds with Löffler *et al.* (Ann Surg Oncol. 2017; 24(6):1650-1657.) and the sample materials obtained during HIPEC used here are identical to those used previously. Values were normalized to 1 at the start of treatment (0 h). A decrease of nCI values signifies cell death of OAW-42 target cells. Graphs show mean ± SD (of 2-6 technical replicates).

**Suppl. Fig. 1 RTCA Pat. 2: Exposure of OAW-42 cells to OCS for 30 minutes at 42 °C *ex vivo***

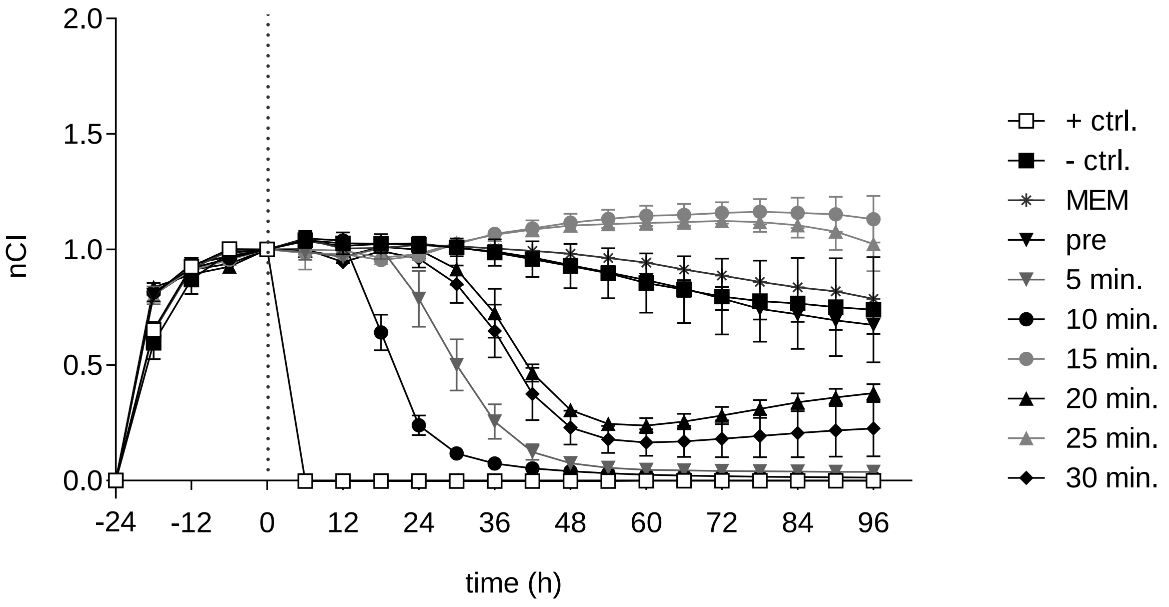


**Suppl. Fig. 2 RTCA Pat. 2: Exposure of OAW-42 cells to OCS for 60 minutes at 42 °C *ex vivo***

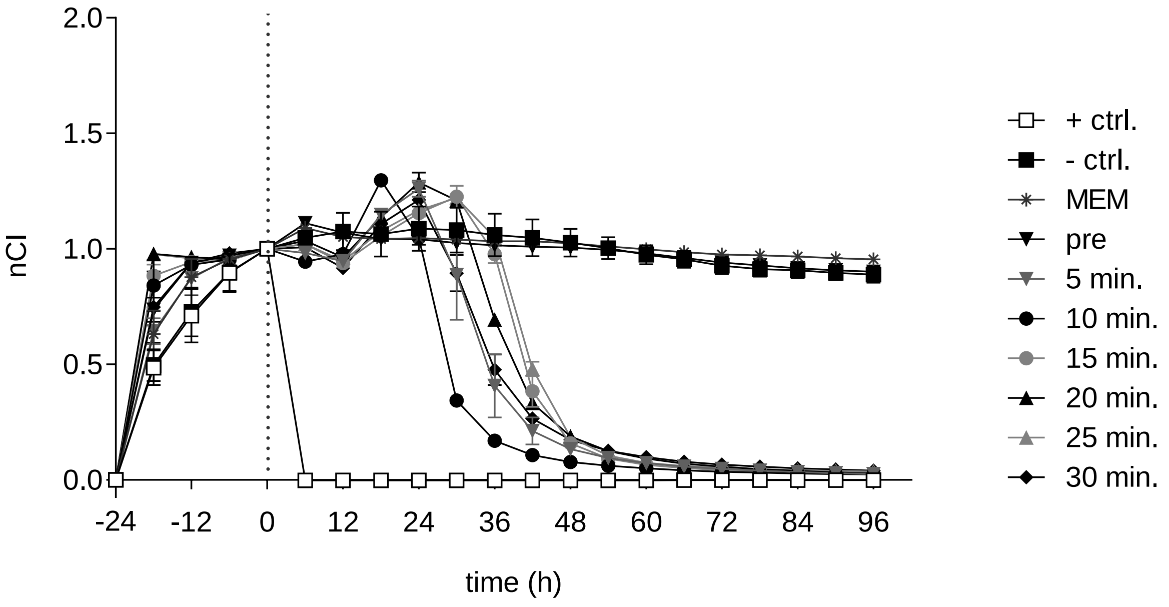

**Suppl. Fig. 3 RTCA Pat. 4: Exposure of OAW-42 cells to OCS for 30 minutes at 42 °C *ex vivo***

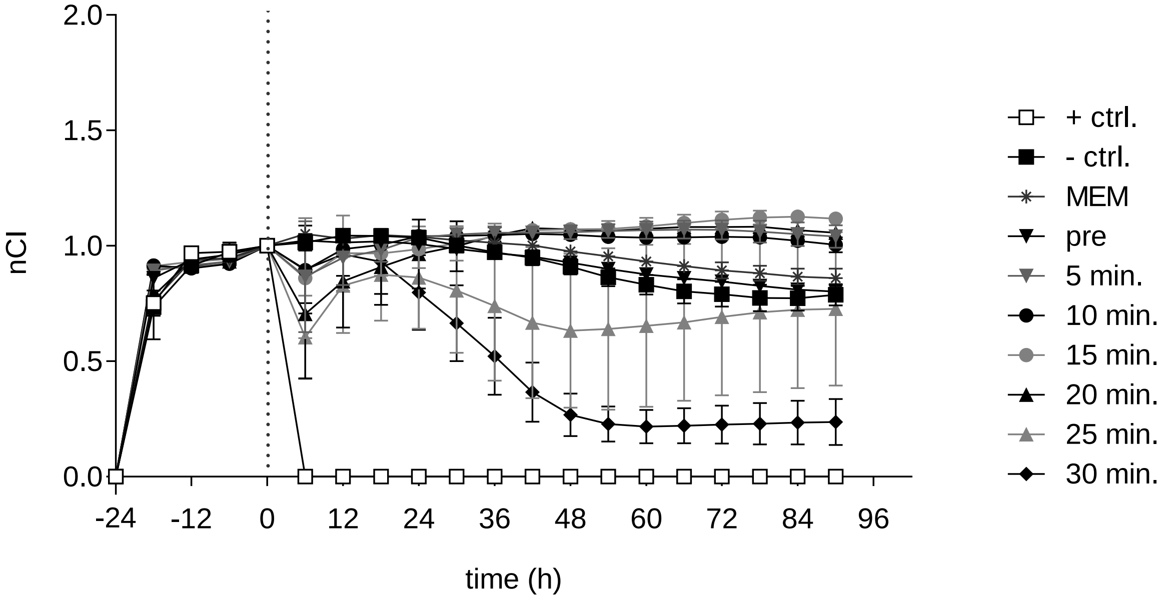


**Suppl. Fig. 4 RTCA Pat. 4: Exposure of OAW-42 cells to OCS for 60 minutes at 42 °C *ex vivo***

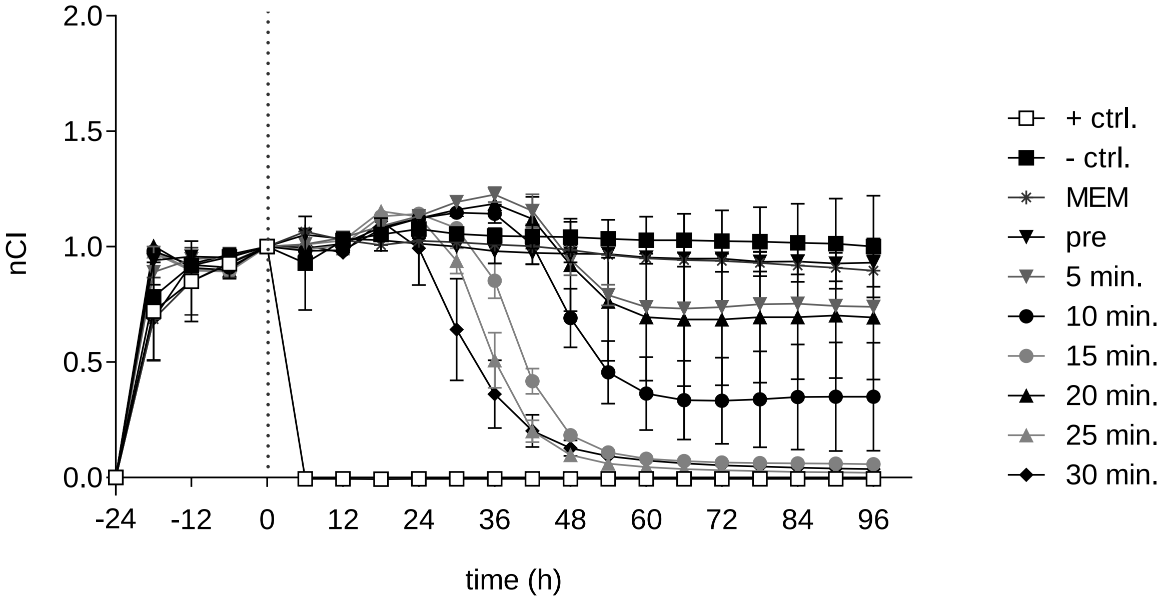

**Suppl. Fig. 5 RTCA Pat. 5: Exposure of OAW-42 cells to OCS for 30 minutes at 42 °C *ex vivo***

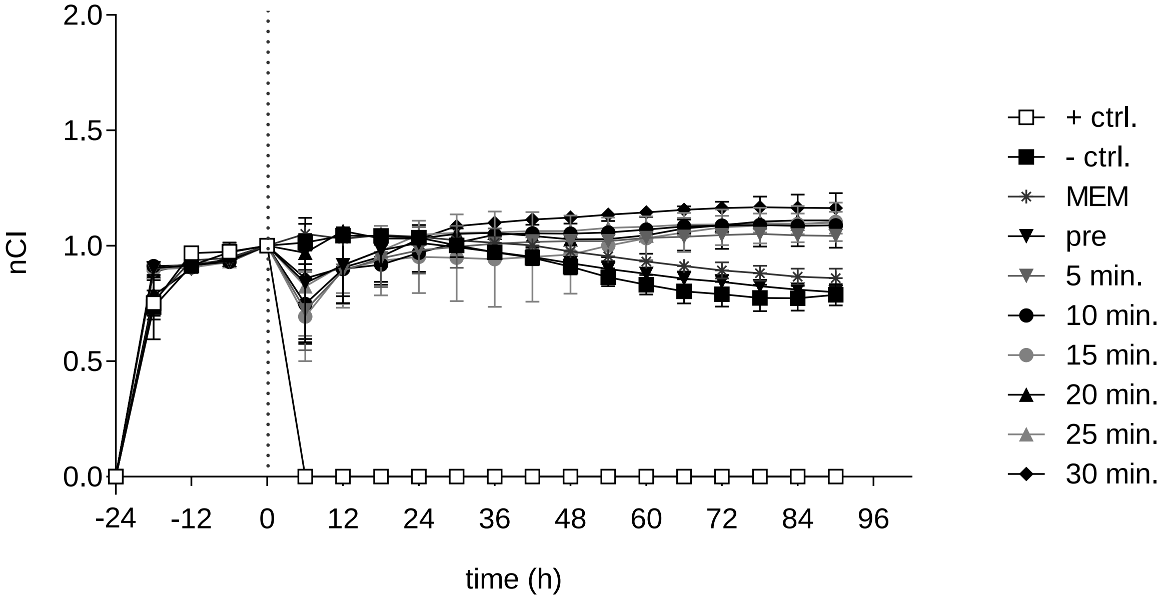


**Suppl. Fig. 6 RTCA Pat. 5: Exposure of OAW-42 cells to OCS for 60 minutes at 42 °C *ex vivo***

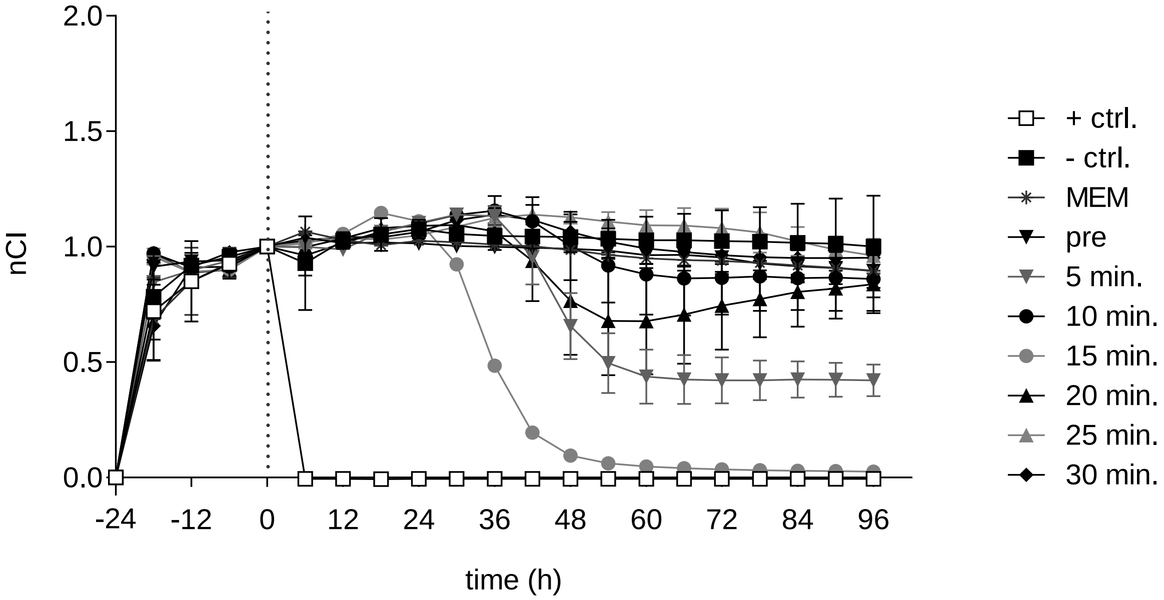

**Suppl. Fig. 7 RTCA Pat. 6: Exposure of OAW-42 cells to OCS for 30 minutes at 42 °C *ex vivo***

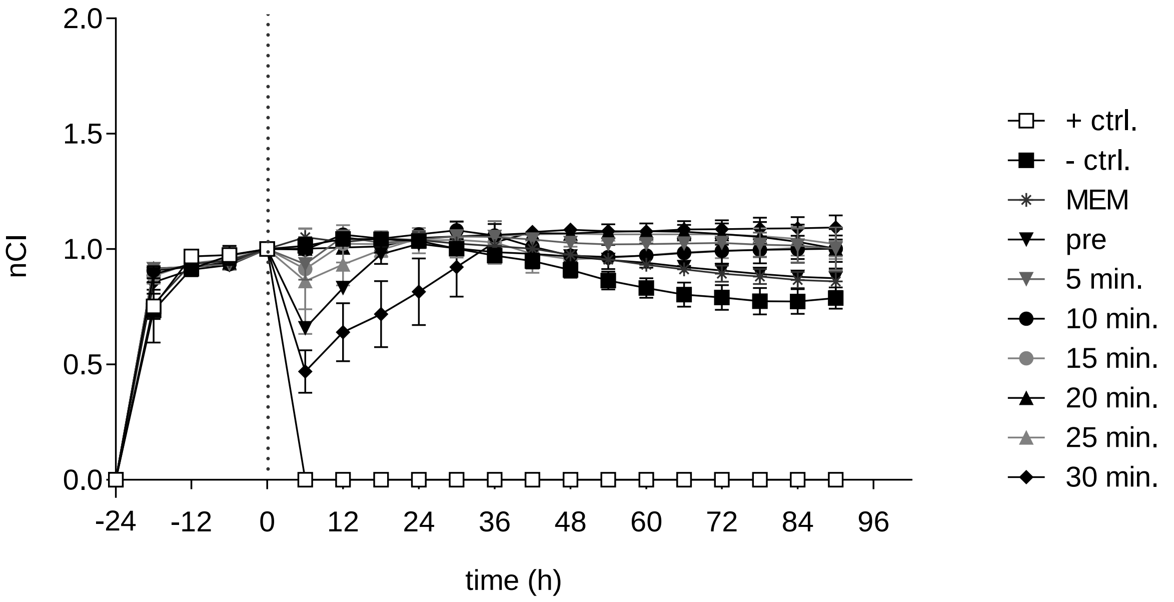


**Suppl. Fig. 8 RTCA Pat. 6: Exposure of OAW-42 cells to OCS for 60 minutes at 42 °C *ex vivo***

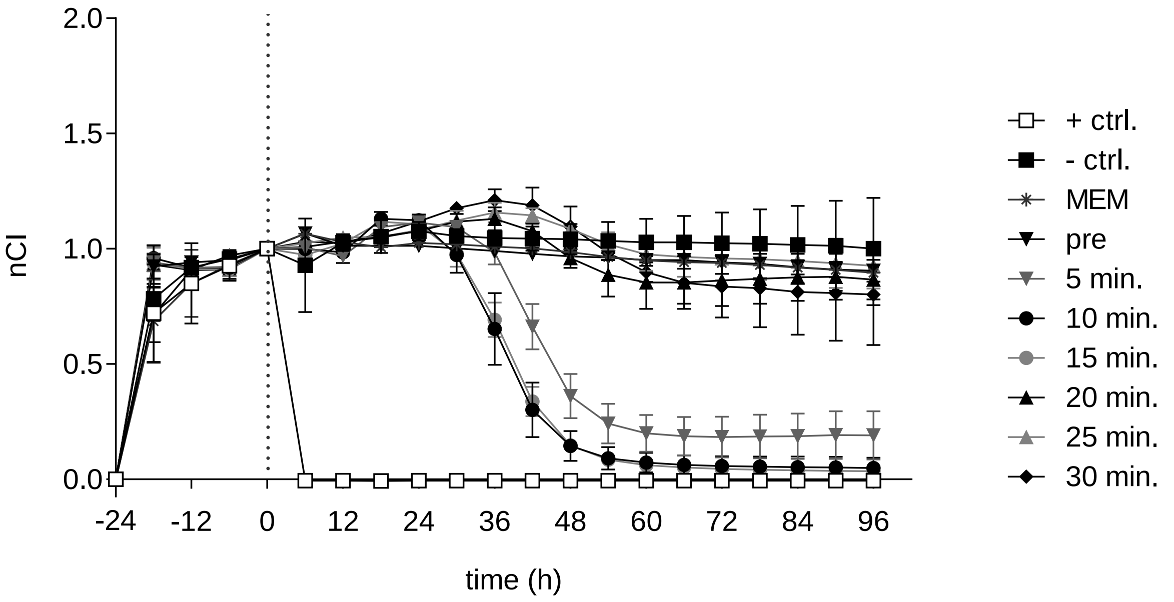

**Suppl. Fig. 9 RTCA Pat. 7: Exposure of OAW-42 cells to OCS for 30 minutes at 42 °C *ex vivo***

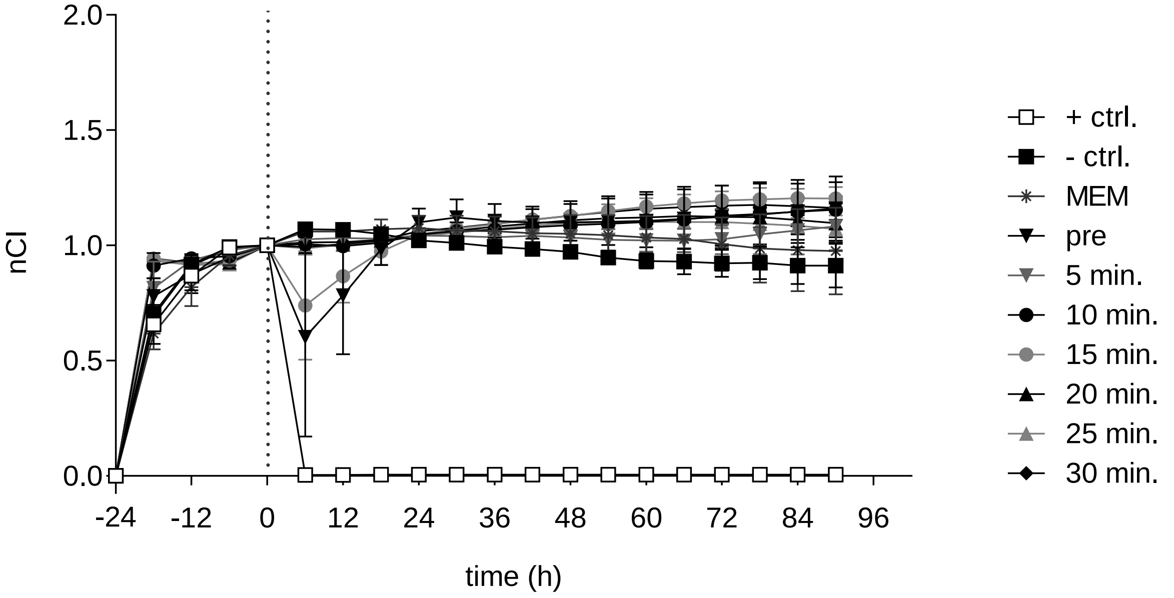


**Suppl. Fig. 10 RTCA Pat. 7: Exposure of OAW-42 cells to OCS for 60 minutes at 42 °C *ex vivo***

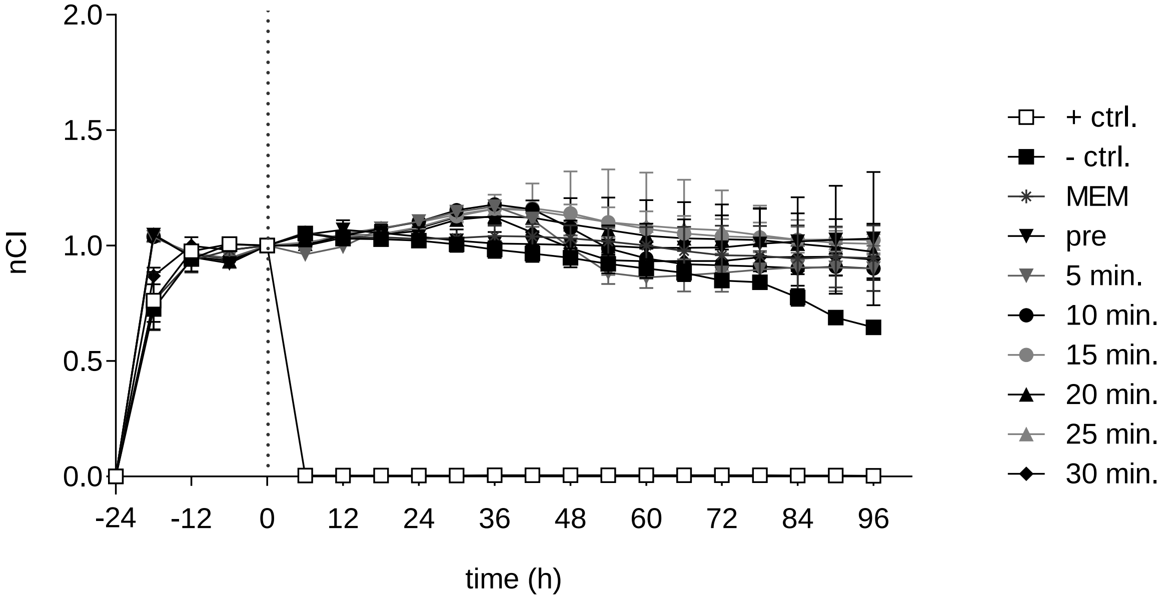

**Suppl. Fig. 11 RTCA Pat. 8: Exposure of OAW-42 cells to OCS for 30 minutes at 42 °C *ex vivo***

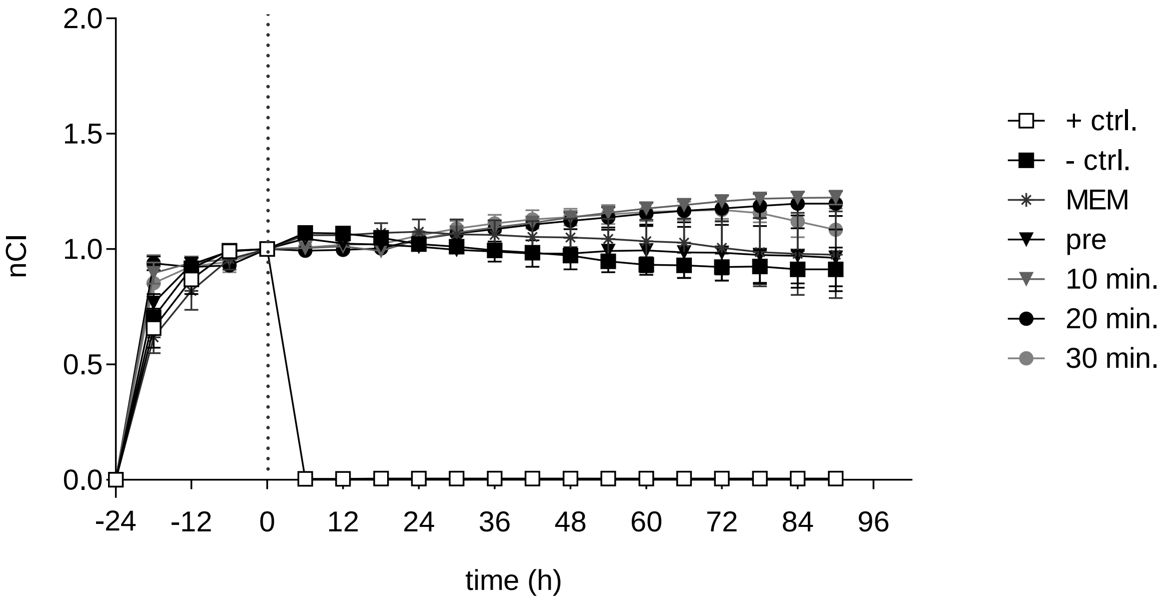


**Suppl. Fig. 12 RTCA Pat. 8: Exposure of OAW-42 cells to OCS for 60 minutes at 42 °C *ex vivo***

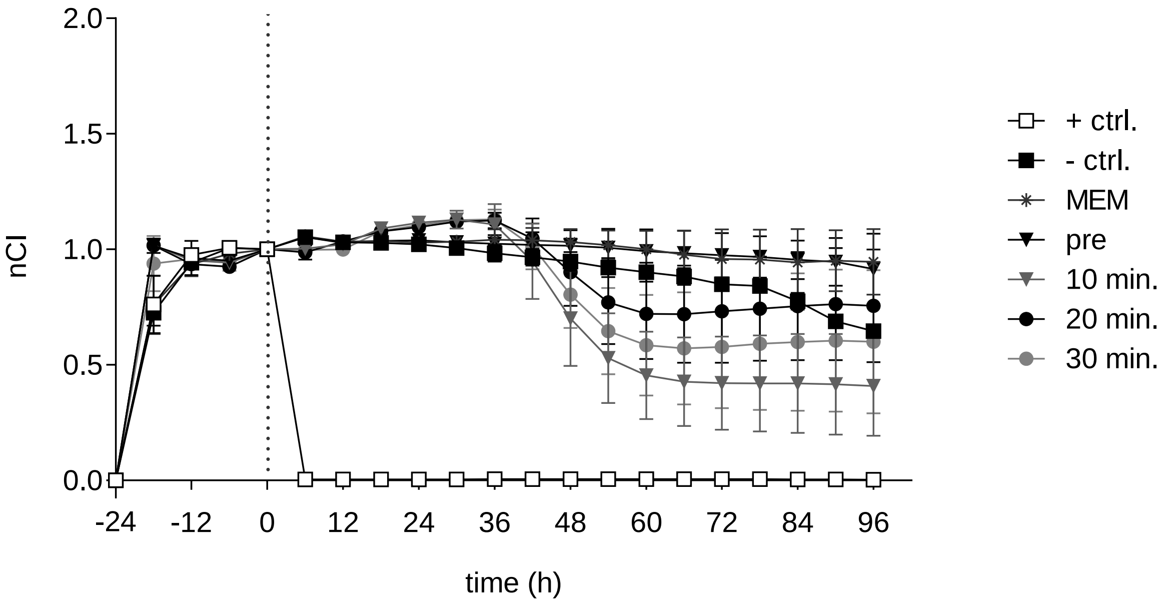

**Suppl. Fig. 13 RTCA Pat. 9: Exposure of OAW-42 cells to OCS for 30 minutes at 42 °C *ex vivo***

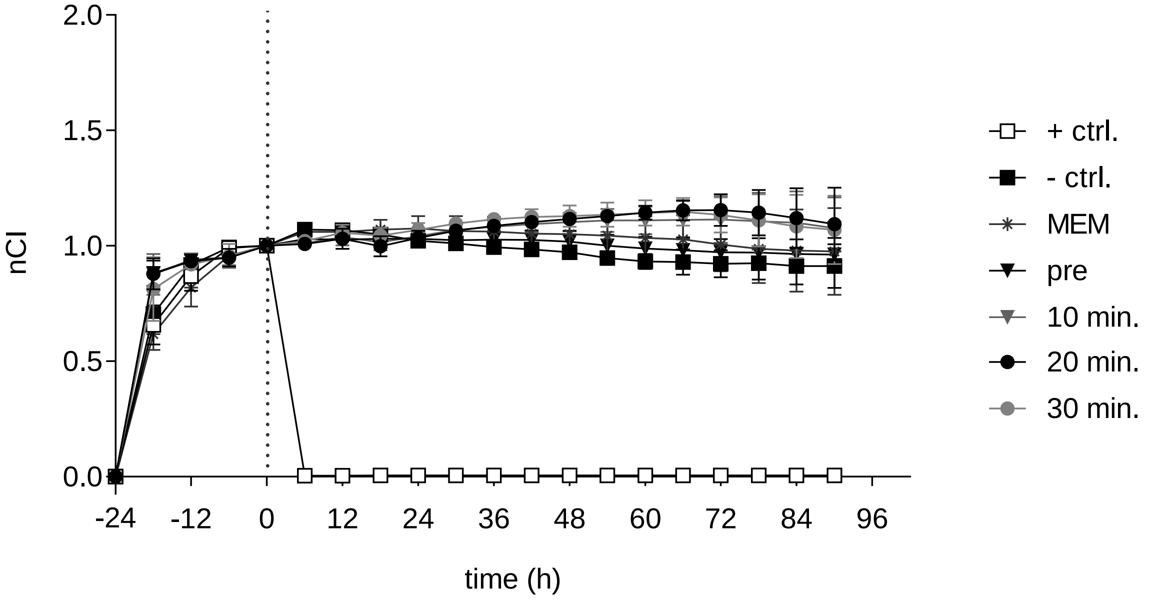


**Suppl. Fig. 14 RTCA Pat. 9: Exposure of OAW-42 cells to OCS for 60 minutes at 42 °C *ex vivo***

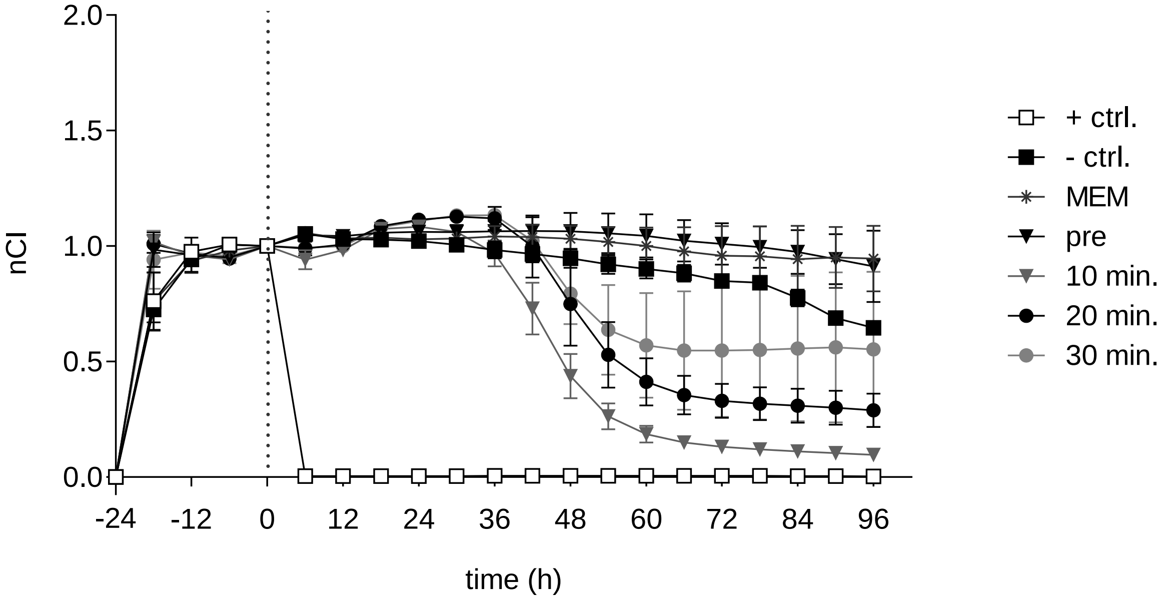

**Supp. Fig. 15 Assessment of the thickness of an OAW-42 cell layer seeded at different densities**

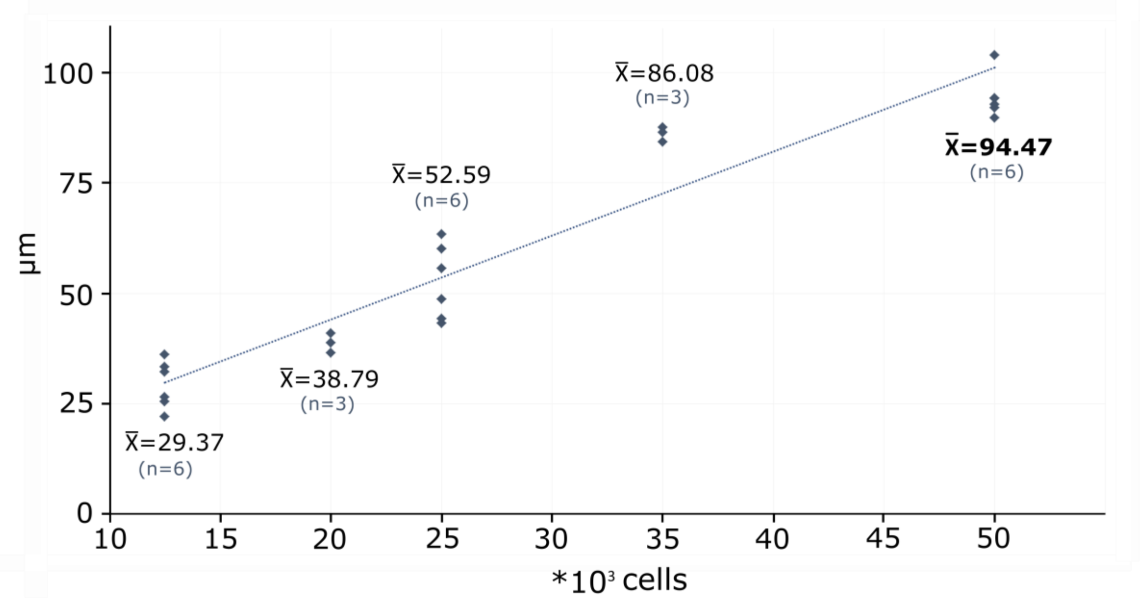


Serial dilutions of OAW-42 cells seeded at different densities of 12.5/ 20.0/ 25.0/ 35.0 and 50.0 x $10^3$ cells/ well (in a 96-well plate) were performed. Cell layer thickness was measured in (n) replicates after 24 hours cell culture, using z-stacks on a Nikon ti eclipse microscope (performed by an unbiased observer) using 10x magnification with the NIS-Elements (Nikon, Tokyo, Japan) or ImageJ software. Experiments were performed twice to obtain measurements at two independent occasions. Arithmetic means ($\bar{x}$) of (n) measurements are given in µm.

**Supp. Fig. 16 Oxaliplatin (OX)-spiked into PDS 60 minutes at 42 °C**

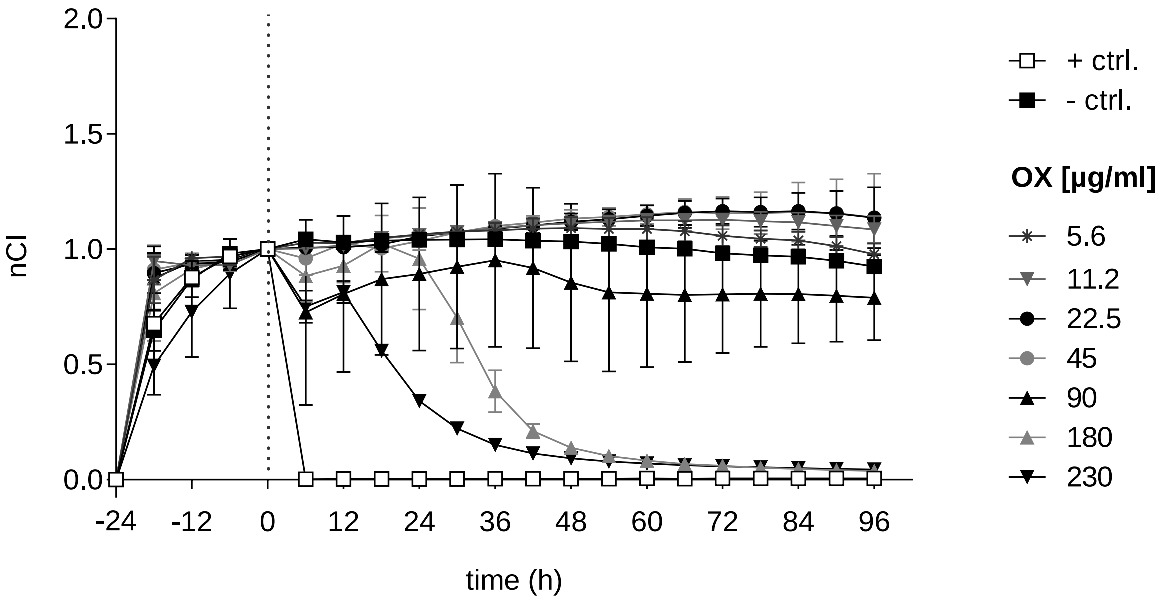


Normalized cell index (nCI) in 6-hour intervals obtained from RTCA impedance measurements of platinum-sensitive OAW-42 cells incubated for 60 minutes at 42 °C with the specified amounts of OX spiked into PDS performed at time point 0 hours (h). (+ ctrl.): Triton; (- ctrl.): Physioneal 40; OX (oxaliplatin); PDS (peritoneal dialysis solution; Physioneal 40); RTCA (real-time cell analysis). Graphs show mean ± SD (of 2-3 technical replicates).

**Supp. Fig. 17 Continuous exposure of OAW-42 cells to oxaliplatin (OX)-spiked into PDS**

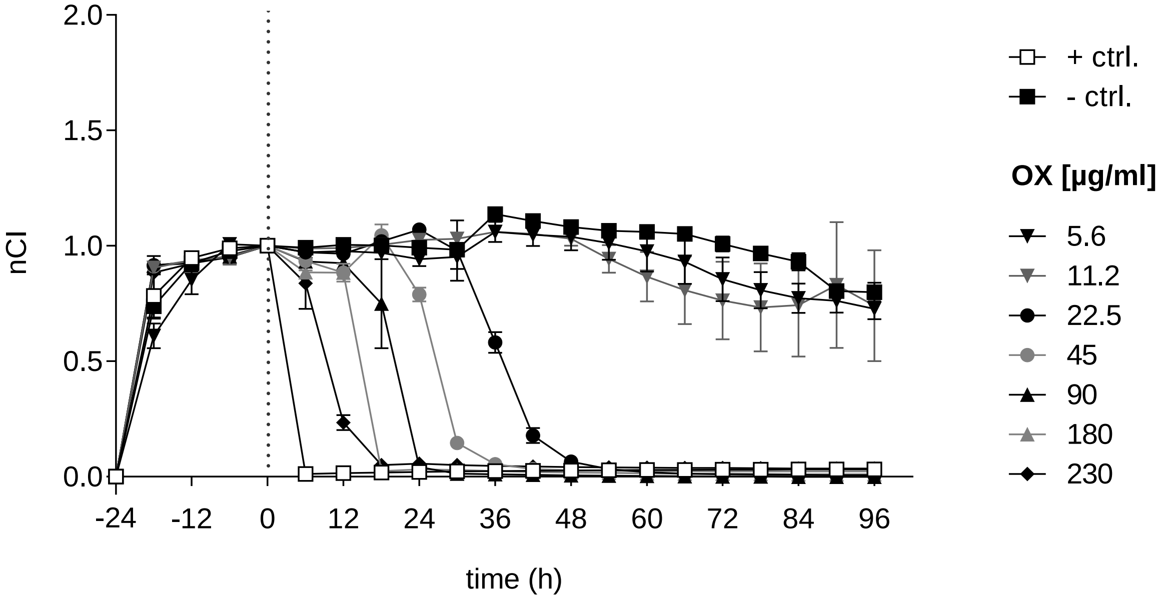


Normalized cell index (nCI) in 6-hour intervals obtained from RTCA impedance measurements of platinum-sensitive OAW-42 cells incubated with the specified concentrations of OX initially spiked into PDS and diluted in MEM (effective end concentrations of OX after dilution in 50 % MEM are given) performed and normalized to 1 at time point 0 hours (h). (+ ctrl.): Triton; (- ctrl.): Physioneal 40. MEM (serum-supplemented cell culture medium); OX (oxaliplatin); PDS (peritoneal dialysis solution; Physioneal 40); RTCA (real-time cell analysis). Graphs show mean ± SD (of 2-6 technical replicates).

**Supp. Fig. 18 Continuous exposure of OAW-42 cells to oxaliplatin (OX)-spiked into D5W**

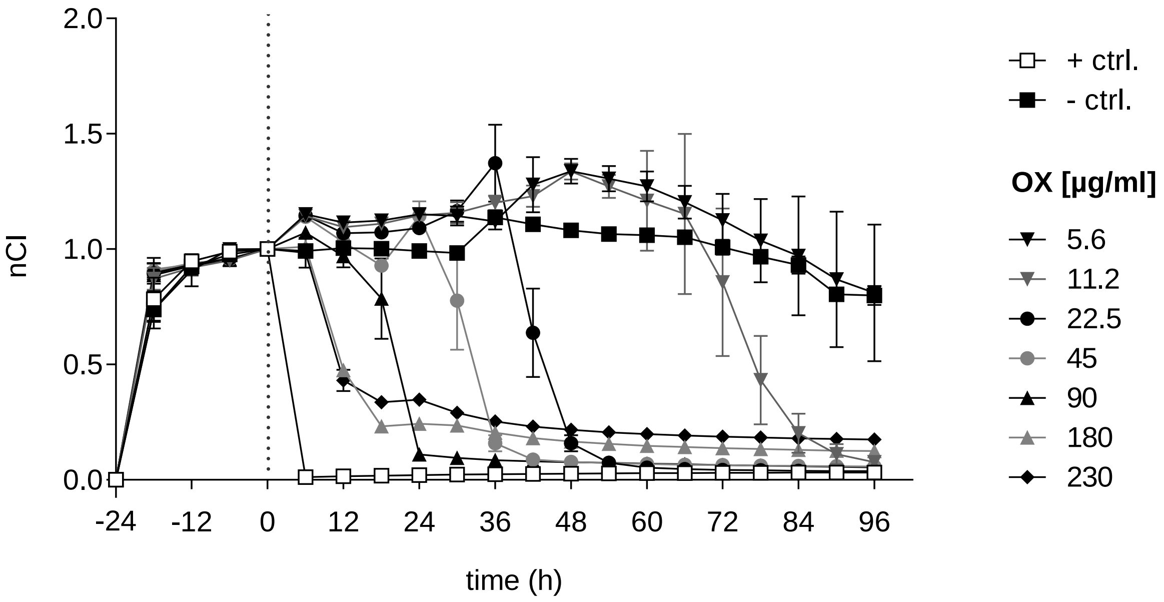


Normalized cell index (nCI) in 6-hour intervals obtained from RTCA impedance measurements of platinum-sensitive OAW-42 cells incubated with the specified concentrations of OX spiked into D5W and diluted in MEM (effective end concentrations of OX after dilution in 50 % MEM are given) performed and normalized to 1 at time point 0 hours (h). (+ ctrl.): Triton; (- ctrl.): D5W: dextrose 5 % in water. MEM (serum-supplemented cell culture medium) OX (oxaliplatin); RTCA (real-time cell analysis). Graphs show mean ± SD (of 2-6 technical replicates).

**Supp. Fig. 19** **End point assays:** ***Cell death of OAW-42/HT29 cells exposed to oxaliplatin (OX)-spiked Ringer´s lactate for 30 minutes at 42 °C***

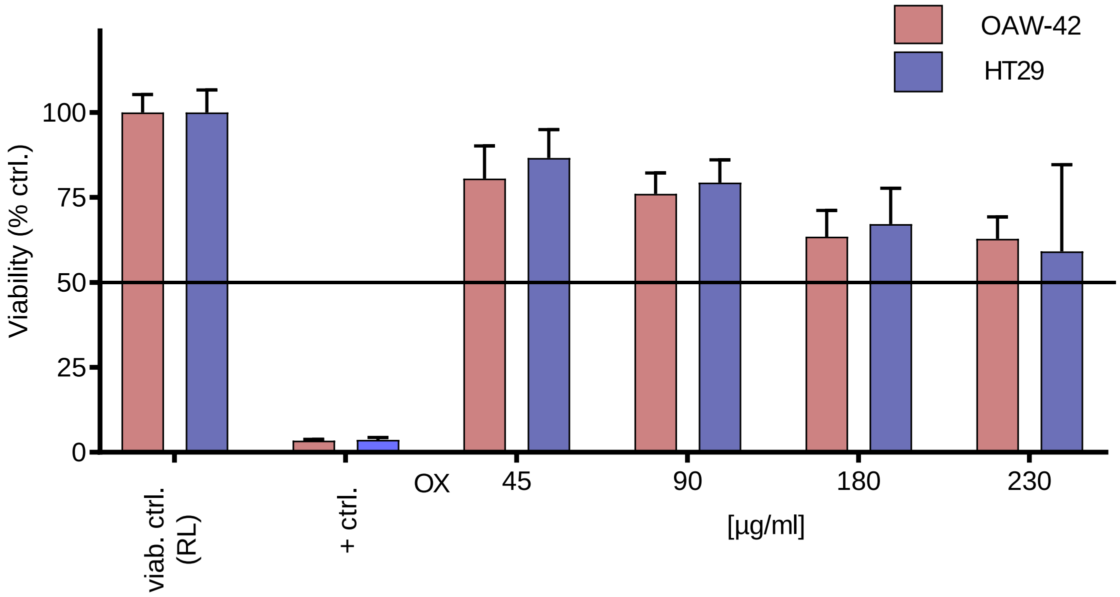


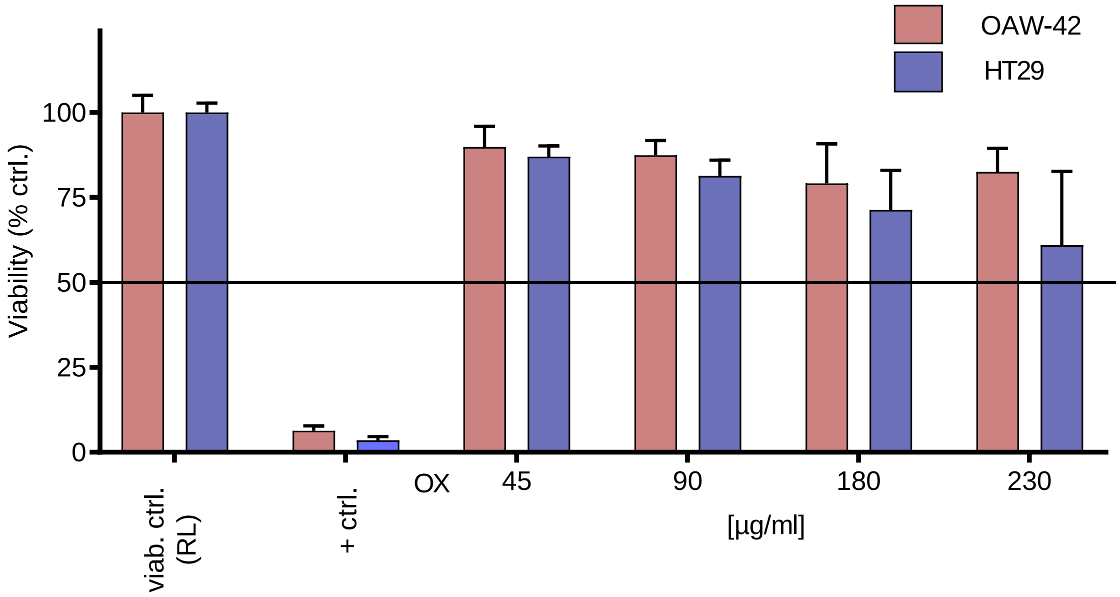


Platinum-sensitive OAW-42 cells and the colon carcinoma cell line HT29 were seeded at a density of 3.15 x $10^5$ cells per well and incubated for 30 minutes at 42 °C with the specified amounts of OX spiked into Ringer´s lactate solution (RL). After OX exposure, cells were washed and subsequently cultured in serum supplemented medium. Afterwards, a fluorometric resazurin-based (CellTiter-Blue®, CTB) (**above**) and a sulforhodamine B (SRB) assay (**lower graph**) was used to determine cell viability. Cells were normalized to cells treated identically with RL. (+ ctrl.): Triton. Graphs show mean ± SD from three independent experiments each performed in triplicates. The $LC_{50}$ threshold is marked with a black line.
